# KRAB zinc finger proteins ZNF587/ZNF417 protect lymphoma cells from replicative stress-induced inflammation

**DOI:** 10.1101/2023.03.08.531722

**Authors:** Filipe Martins, Olga Rosspopoff, Joana Carlevaro-Fita, Romain Forey, Sandra Offner, Evarist Planet, Cyril Pulver, HuiSong Pak, Florian Huber, Justine Michaux, Michal Bassani-Sternberg, Priscilla Turelli, Didier Trono

## Abstract

Heterochromatin loss and genetic instability enhance cancer progression by favoring clonal diversity, yet uncontrolled replicative stress can lead to mitotic catastrophe and inflammatory responses promoting immune rejection. KRAB-containing zinc finger proteins (KZFPs) are epigenetic modulators, which for many control heterochromatin at transposable element (TE)-embedded regulatory sequences. We identified a cluster of 18 KZFPs associated with poor prognosis in diffuse large B cell lymphoma (DLBCL). We found their upregulation to correlate with increased copy number alterations and suppression of immune responses in tumor samples. Upon depleting two that target evolutionarily recent TEs, the primate-specific ZNF587 and ZNF417 paralogs, the proliferation of DLBCL cell lines was drastically impaired and replicative stress abruptly induced with marked alterations of the chromatin landscape and multiplication of DNA replication origins. Furthermore, *ZNF587/417* knockdown upregulated interferon/inflammatory-related genes through activation of the cGAS-STING DNA sensing pathway, augmented the susceptibility of tumor cells to macrophage-mediated phagocytosis, and modified their immunogenicity through an increased surface expression of HLA-I and reshuffling of their immunopeptidome. ZNF587 and ZNF417 are thus pro-oncogenic factors allowing for higher degrees of genetic instability through attenuation of replicative stress and secondary inflammation, an influence that likely facilitates the clonal expansion, diversification, and immune evasion of cancer cells.

## INTRODUCTION

Diffuse large B cell lymphoma (DLBCL), the most common type of lymphoid malignancy^1,2^, is plagued by relapse or refractoriness to first-line therapy in nearly fifty percent of cases^3^. DLBCL arises from the neoplastic transformation of mature B cells during the final steps of their differentiation into antibody-secreting plasma cells or memory B cells^4^. This program is normally under tight control by epigenetic mechanisms ensuring the coordinated activation of specific gene expression networks^5^, but it can be dysregulated by aging^6^, mutations in critical genes^7^ and other environmental influences^8^, leading to lymphomagenesis. Three major molecular subtypes of DLBCL with prognostic relevance^9,10^ have been delineated based on similarities between the transcriptome of these tumors and their putative cell of origin (COO). Germinal center B cell (GCB) and activated B cell (ABC) DLBCLs are thought to derive from B cells blocked at the centroblast and plasmablast stages, respectively, while the transcriptome of so-called “unclassifiable” DLBCL does not evoke any of these two differentiation stages.

Genomic instability is a primary driver of the clonal evolution and diversification of cancer cells^11,12^. It frequently results from errors occurring during genome replication, which starts with the assembly of pre-replicative complexes (pre-RCs) during the G1 phase of the cell cycle, an event known as origin licensing^13^. During early S phase, pre-RCs recruit the DNA polymerase and elongation factors to form the replication fork machinery. Subsets of licensed replication origins then become activated in a sequential and spatially regulated manner^14^, allowing the bidirectional progression of replication forks across the genome. Conducting a proper replication program in the presence of short cell cycling^15,16^, intercurrent mutagenic events^17^ and exacerbated oncogene-induced transcriptional activity^18^ is challenging, and accordingly DLBCL cells are prone to replicative stress (RS), characterized by the stalling and potential collapse of replication forks leading to genomic instability^19^.

Increasing evidence suggests a central role for chromatin compaction in the selection and timing of activation of DNA replication origins^20^. Heterochromatin regions are known to be late-replicating, a phenomenon attributed to their high content in repetitive DNA sequences, a major source of RS as they provide hotspots for homologous recombination, secondary DNA structures, and the formation of RNA:DNA hybrids^21–25^. Pericentromeric regions and telomeres, which are enriched for satellite and simple repeats, are the most heterochromatin-dense regions of the genome. Nevertheless, some 4.5 million transposable elements (TEs) sprinkled over the rest of the human genome also contribute significantly to its heterochromatin landscape. Most human TEs are retroelements belonging to the LTR (long terminal repeat)-containing endogenous retroviruses (ERV), long and short interspersed nuclear elements (LINE and SINE, respectively) or SVA (SINE-VNTR-Alu-like) families, which replicate by a “copy-and-paste” mechanism through reverse transcription of an RNA intermediate and integration of its DNA copy elsewhere in the genome. TEs are motors of genome evolution^26^, and a major source of *cis*-acting regulatory sequences that shape gene regulatory networks at play in broad aspects of human biology^27–29^. The disruptive potential of TEs has led to the evolutionary selection of many protective mechanisms, whether to prevent transposition during genome reprogramming in the germline and early embryogenesis or to control the transcriptional impact of TE-embedded regulatory sequences. Repression of TE loci via DNA methylation and heterochromatin formation through methylation of lysine 9 of histone 3 (H3K9me3), stand prominently amongst these mechanisms, and Krüppel-associated box (KRAB) zinc finger proteins (KZFPs) amongst their mediators^30^.

KZFPs constitute the largest family of transcription factors (TFs) encoded by higher vertebrates, with nearly 400 members in human alone, more than a third of them primate-restricted and the products of recent gene duplication events^31,32^. KZFPs bind DNA in a sequence-specific manner through their C-terminal arrays of zinc fingers, and a large majority of human KZFPs have TE-embedded sequences as primary genomic targets^31^. Most act as transcriptional repressors via the KRAB domain-mediated recruitment of KAP1 (KRAB-associated protein 1, also known as tripartite motif protein 28 pr TRIM28)^33–35^, which serves as a scaffold for the assembly of a heterochromatin-inducing complex comprising nucleosome remodeling, histone deacetylase, and histone and DNA methyltransferase activities^33^. Rather than just another silencing mechanism aimed at preventing TE mobilization, KZFPs are increasingly recognized as facilitating the domestication of TEs regulatory potential^31,36^. Indeed, KZFPs not only repress TEs immediately after genomic activation but also corral their transcriptional influences at various stages of development and in adult tissues^37–41^. Thus, KZFPs both minimize the genotoxic potential of TEs and partner up with their genomic targets to shape lineage- and species-specific transcription regulatory networks^42,43^.

KZFPs have been proposed to exert collectively tumor suppressive functions across many tumor types^44^. However, individual family members have been linked to carcinogenesis through the deregulation of genes implicated in cell cycle progression^45^, apoptosis^46,47^, epithelial to mesenchymal transition (EMT)^48^, chemoresistance^49^, and metastasis^44,50^, and have been proposed to promote disease in lymphocytic leukemia^51^, lung^52,53^, colorectal^54^, and ovarian cancer^55^. In addition, their main cofactor KAP1 functions as an E3-ubiquitin ligase capable of inactivating p53^56^, and the KAP1-recruited SETDB1 methyltransferase was recently found to foster immune evasion in melanoma and lung cancer by repressing TE transcription, a source of dsDNA/dsRNA species sensed by intracellular innate immunity pathways and of endogenous viral peptides with immunogenic properties^57,58^.

Here, we took advantage of a large dataset of DLBCL patients with extensive clinical and molecular documentation^59^ to explore the potential role of KZFPs in this hematological malignancy. This led us to discover that ZNF587 and ZNF417, a pair of primate-specific KZFP paralogs targeting evolutionarily recent TEs and so far mostly known for their involvement in human brain development and the control of neuronal gene expression, protect DLBCL cells from excessive replicative stress associated with genome-wide alterations of the heterochromatin landscape, thus safeguarding these cells from cell-intrinsic inflammatory responses.

## RESULTS

### A DLBCL transcriptome-wide association study reveals KZFPs as predictors of poor prognosis

To probe the potential involvement of KZFPs in DLBCL pathogenesis, we looked for a correlation between their levels of expression and disease outcome. For this, we used published RNA-seq data from 633 fresh-frozen paraffin-embedded DLBCL samples collected before the initiation of rituximab-based therapy^59^. We conducted a univariate Cox proportional hazards regression analysis on the transcriptome of these samples to identify genes, the expression of which was either positively or negatively correlated with survival. Out of 23’560 expressed genes (see methods), 1’530 (6.5%) were found to be significantly associated with either a better (n=950, 62.1%) or a worse (n=580, 37.9%) outcome (Fig. 1a). KZFP genes were significantly enriched in prognosis-associated genes with 41 out of 395 (10.4%) expressed KZFPs associated with survival (1’530/23’560 vs. 41/395; Fisher’s exact test, *p* = 0.0038), 39 of them negatively. This stood in sharp contrast with the observed distribution of hazards ratios (HRs) for all other TFs or remaining protein-coding genes, where good-prognosis hits dominated (Fig. 1b). Corroborating our results, a recent analysis of the same cohort found 12 of these 39 KZFP genes to be part of a KZFP-enriched poor-prognosis gene expression signature^60^.

**Fig. 1.**
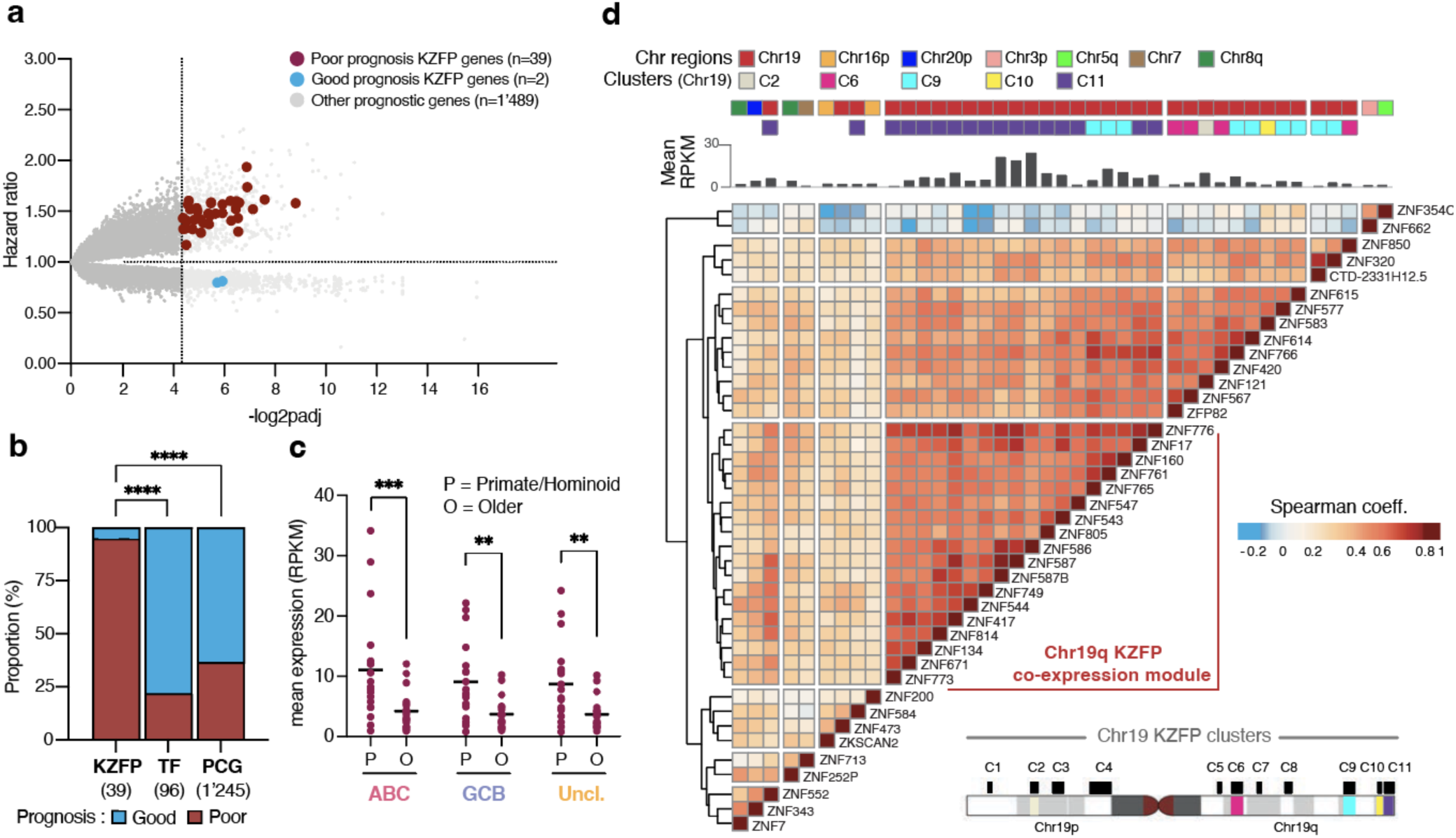
A *KZFP* gene cluster as poor prognosis predictor in DLBCL. (a) Volcano plot of hazard ratios (HRs) issued from univariate Cox regression analysis correlating individual gene expression with patient survival outcome are plotted against significance (n=612 DLBCL RNA-seq with available survival data). The vertical dashed line indicates the threshold of significance (FDR < 0.05). The horizontal dashed line separates genes associated with an increased (>1) and reduced risk (<1) of death. (b) Stacked bar plots showing the proportion of gene hits associated with good (blue) or poor prognosis (dark red). Genes were hierarchically categorized as protein-coding KZFPs, other TFs, and other protein-coding genes (PCGs). Statistics: Two-tailed Fisher’s Exact Test. (c) Dot plot showing the mean expression in reads per kilobase of exon per million reads mapped (RPKM) of the 19 primate/hominoid-specific (P) and 20 evolutionary older (O) protein-coding KZFP gene hits correlated with survival across ABC (n=256), GCB (n=281) and Unclassifiable (n=96) DLBCL samples. Statistics: Two-sided Mann–Whitney U-test. (d) Hierarchical clustering and heatmap showing the spearman correlation coefficients of pairwise comparison between the 41 KZFP gene hits based on their gene expression across 633 DLBCL samples – Euclidean distances and complete-linkage clustering method. The bar graph on top of the heatmap shows the mean expression in RPKM and the upper annotation track shows the chromosomal location. The lower right ideogram displays KZFP gene hits positioning onto previously described Chr19q KZFP clusters^43^. Statistical significance is indicated by the asterisks as follows: *, *p* < 0.05; **, *p* < 0.01; ***, *p* < 0.001; ****, *p* < 0.0001.

Among the 41 *KZFP* gene hits of our screen, 39 were protein-coding, 19 of them primate-specific and expressed at higher levels than their evolutionary older counterparts in all disease subtypes (Fig. 1c)^31^, suggesting a possible (COO-independent) pro-oncogenic role for these proteins despite their young evolutionary age. *ZNF662* and *ZNF354C*, the two better-prognosis *KZFP* genes identified by our analysis, reside on chromosomes 3 and 5, respectively. Interestingly, the long arm of chromosome 19 (Chr19q) harbors 32 of the 39 (82.1%) poor prognosis *KZFP* genes, a significant overrepresentation of this genomic region as the entire chromosome 19 only encodes slightly more than half of all KZFPs (231/395, 58.5%; Fisher’s exact test, *p* = 0.0034)^61,62^. To evaluate if these *KZFP* genes were part of a single gene expression module, we asked which were systematically co-expressed in DLBCL samples. This led us to identify a subset of 18 *KZFPs* displaying a strong co-expression pattern. Fifteen of these were located within the C11 KZFP gene cluster located on the sub-telomeric region of chromosome 19 (chr19q13.43) and the other 3 within the nearby C9 cluster (Fig. 1d)^43^. This prompted us to characterize the DLBCL subset overexpressing these coregulated unfavorable *KZFP* genes.

### A *KZFP* gene cluster defines a DLBCL subset with cell-autonomous growth features

Through unsupervised hierarchical clustering of DLBCL samples based on the expression of these 18 co-expressed *KZFP* genes, we could delineate a subset encompassing about a quarter of all DLBCL samples (n=145, 23%, Fig. 2a), to which we will refer to hereafter as KZFP^High^ DLBCLs. To characterize differences between these samples and their KZFP^Low^ counterparts, we conducted a gene set enrichment analysis (GSEA)^63,64^ for pathways catalogued in the Hallmark gene sets^65^. KZFP^High^ samples were characterized by a proliferative and highly metabolic state, as suggested by the upregulation of genes involved in cell cycle regulation, DNA repair, mitotic spindle assembly, oxidative phosphorylation and Myc response. Conversely, KZFP^Low^ samples were characterized by the activation of interferon (IFN)/inflammatory response and EMT (epithelial-to-mesenchymal transition) gene sets (Fig. 2b).

**Fig. 2:**
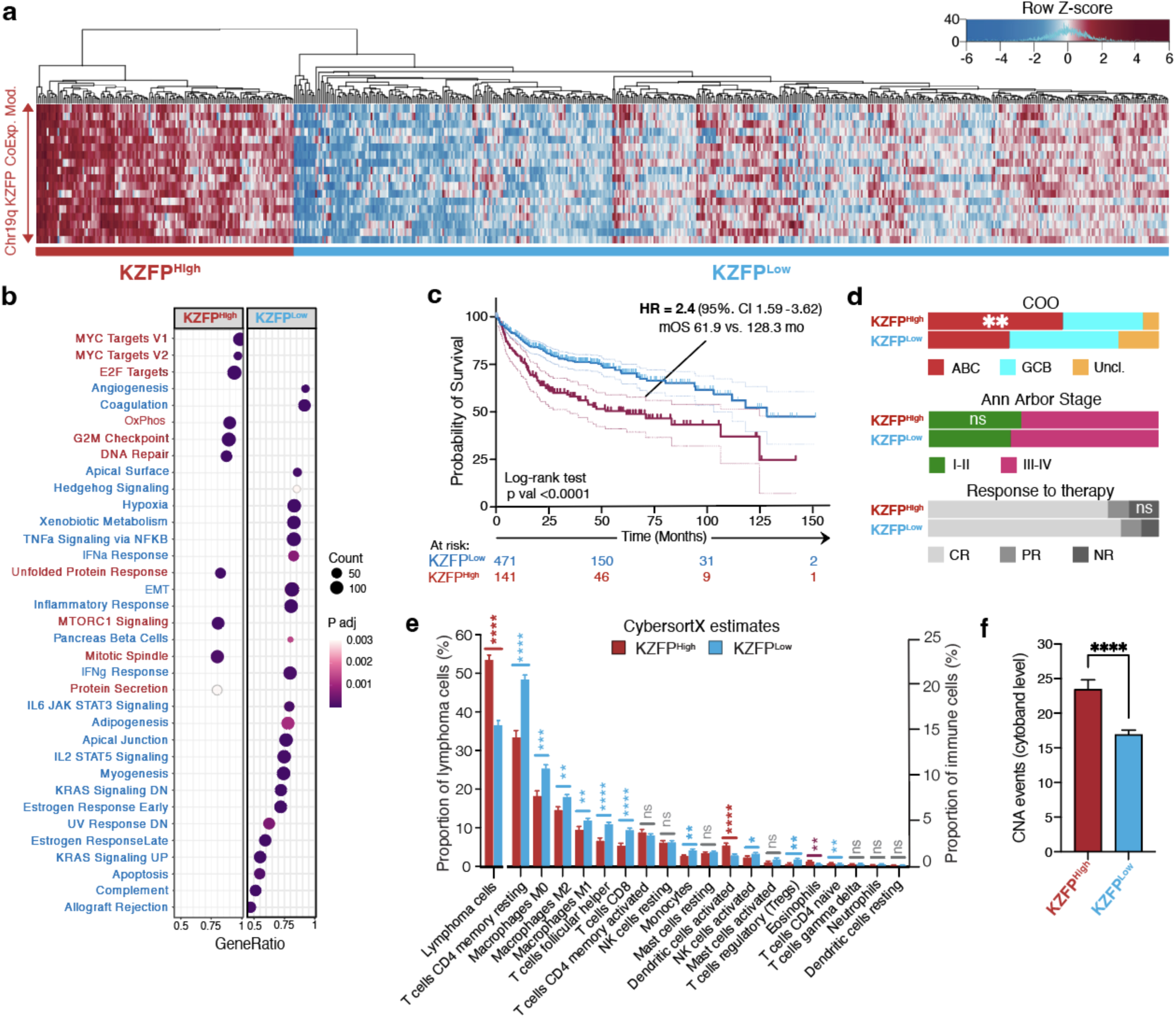
KZFP^High^ DLBCLs display cell-autonomous growth features. (a) Heatmap of the unsupervised hierarchical clustering analysis of 18 KZFP genes expression of the Chr19q co-expression module across 633 DLBCL samples – Euclidean distances and Ward’s agglomeration method. (b) Dot plots of enriched Hallmark gene sets in KZFP^High^ vs. KZFP^Low^ DLBCLs. The X-axis represents the ratio of differentially expressed genes (DEGs) in these groups over the total number of genes of each pathways (gene ratio). Dot size represent the number of DEGs in each gene set (count) and color scale represents the False Discovery Rate (FDR <0.05). (c) Kaplan-Meier survival curves of patients displaying a KZFP^High^ vs. KZFP^Low^ DLBCL signature as defined in (a). Statistics: Log-rank test. (d) Stacked bar plots showing the proportion of the 3 different COO subtypes, the Ann Arbor stages at diagnosis, and the response to initial rituximab-based standard regimen amongst the patients belonging to each KZFP-defined group. CR = complete response; PR = partial response; NR = no response. Statistics: Two-tailed Fisher’s Exact Test. (e) Bar plot showing the differences in the mean proportion (+/− SEM) of lymphoma cells (left) and immune cell infiltrate (right) between KZFP^Low^ and KZFP^High^ DLBCLs using CIBERSORTx RNA-seq bulk deconvolution. Statistics: Two-sided Mann–Whitney U-test with Benjamini &Yekutieli correction. (f) Bar plots showing the mean number (+/−SEM) of copy number alterations (CNA) detected by WES conducted on the same samples included in the transcriptomic analysis.

Using the metadata available for this patient cohort^59^, we determined that individuals presenting with KZFP^High^ DLBCL had survival rates twice shorter than those with a KZFP^Low^ profile (Fig. 2c). KZFP^High^ patients were slightly older (65 +/−13.2 vs 59 +/−16.2 years) and presented more frequently DLBCLs of the ABC subtype (47.2% vs. 34.7%) (Fig. 2d), but were diagnosed at similar disease stages (Fig. 2d). To assess the independence of these KZFP risk groups and the outcome predicted by age, stage at diagnosis and COO, we conducted a multivariate Cox regression analysis, which confirmed that KZFP risk groups remained significantly associated with survival (HR= 1.31, *p* = 0.04). Intriguingly, KZFP^High^ DLBCLs did not appear to present a lower response rate to first-line rituximab-based regimens (Fig. 2d), suggesting that patients in this group were instead more inclined to relapse after first achieving remission. Although the data was not available to confirm directly this point, a Kaplan Meier analysis revealed that the steepest decrease in the survival rate of KZFP^High^ patients occurred during the first 2.5 years following diagnosis (Fig. 2c), which is the most at-risk period for DLBCL relapse^66^. Furthermore, analysis of the HR by stratifying KZFP^High/Low^ groups showed that patients in the KZFP^High^ group had a higher hazard of death peaking in the first 6 months after diagnosis, which remained higher than their KZFP^Low^ counterparts up to 2-3 years, supporting the suspicion of higher incidence of relapses in the former group (Fig. S1a).

This led us to hypothesize that *KZFP* expression may contribute to disease relapse not by causing primary chemo-resistance of the bulk population of lymphoma cells, but rather by contributing to their clonal diversity, thus facilitating the emergence of treatment-resistant subclones, and by fostering the immune evasion of residual disease. To test this hypothesis, we applied a bulk RNA-seq deconvolution method^67^ to DLBCL samples as a proxy for comparing the tumor microenvironment (TME) cell compositions of KZFP^High^ and KZFP^Low^ subtypes. This revealed a significant depletion of both innate and adaptative immune cells in KZFP^High^ compared with KZFP^Low^ DLBCL samples (Fig. 2e). This drop in immune gene signatures concerned T cell subtypes, activated Natural Killer (NK) cells, and monocyte-derived cells, with the exception of activated dendritic cells and eosinophils. Thus, KZFP^High^ DLBCLs triggered weaker immune responses than their KZFP^Low^ counterparts. These apparent differences in KZFP^High/Low^ TME were further supported by the analysis of the mutational pattern of the KZFP-defined DLBCL groups, with KZFP^High^ samples displaying significant enrichment for a recently defined high-risk molecular gene signature linked to *MYD88* and *CD79B* mutations, together with an increased prevalence of *BCL2* and *CDKN2A* alterations (Fig. S1b)^68,69^. This mutational signature, known as the MCD subtype, has been shown to promote DLBCLs immune evasion by reducing antigen presentation and NK cell activation amongst other mechanisms^70^. In addition, the analysis of whole-exome sequencing (WES) data issued from the same samples indicated a higher number of copy number alterations (CNAs) amongst KZFP^High^ DLBCLs, compatible with greater clonal diversity (Fig. 2f).

Taken together, these results indicate that KZFP^High^ DLBCLs are characterized by a proliferative, genetically unstable and cell-autonomous phenotype, as well as an increased frequency of genetic changes associated with aggressive forms of the disease.

### ZNF587/417 depletion impairs lymphoma cell growth and viability

The most highly expressed poor prognosis-associated *KZFPs* across DLBCL samples, *ZNF586, ZNF587B,* and *ZNF587*, are located in a 200kb region of the Chr19q13.43 cytoband in the vicinity of other poor-prognosis *KZFPs* (*ZNF417, ZNF814, ZNF776, ZNF671,* and *ZNF552*). As a first step to investigate the impact of these genes on DLBCL homeostasis, we performed proliferation assays in U2932 and OCI-Ly7 cells, respectively representative of ABC and GCB DLBCL, transduced with lentiviral vectors expressing small hairpin RNAs (shRNAs) against these 3 KZFPs and their respective paralogs, or a scrambled sequence as a negative control (Fig. S2a). *ZNF587B*/*ZNF814* and *ZNF417/ZNF587*, two pairs of paralogs with respectively 70% and 98% of sequence identity, could be simultaneously repressed with a single shRNA targeting shared homologous regions in both paralog pairs. Although highly expressed in these cells (Fig. S2b), *ZNF587B/814* could be knocked down without affecting the growth of the U2932 ABC or OCI-Ly7 GCB cell lines (Fig. 3a). In contrast, ZNF586 depletion stopped the proliferation of U2932 but was less efficient in decreasing OCI-Ly7 cell growth, whereas *ZNF587/417* knockdown (KD) triggered abrupt growth arrest in both settings (Fig. 3a). This result was confirmed with a second shRNA targeting *ZNF587/417* with similar KD efficiency (Fig. S2a/c) and in two additional cells lines, with growth arrest induced in both HBL1 (ABC) and SUDHL4 (GCB) cells, for the latter with a marked reduction in cell viability (Fig. 3b). Most importantly, knockdown of *ZNF587* and *ZNF417* did not preclude the proliferation of normal diploid primary foreskin fibroblasts (Fig. S2d), as similarly observed in human embryonic stem cells^71^, suggesting that dependence on these two KZFPs is somewhat restricted to cancer cells.

**Fig. 3:**
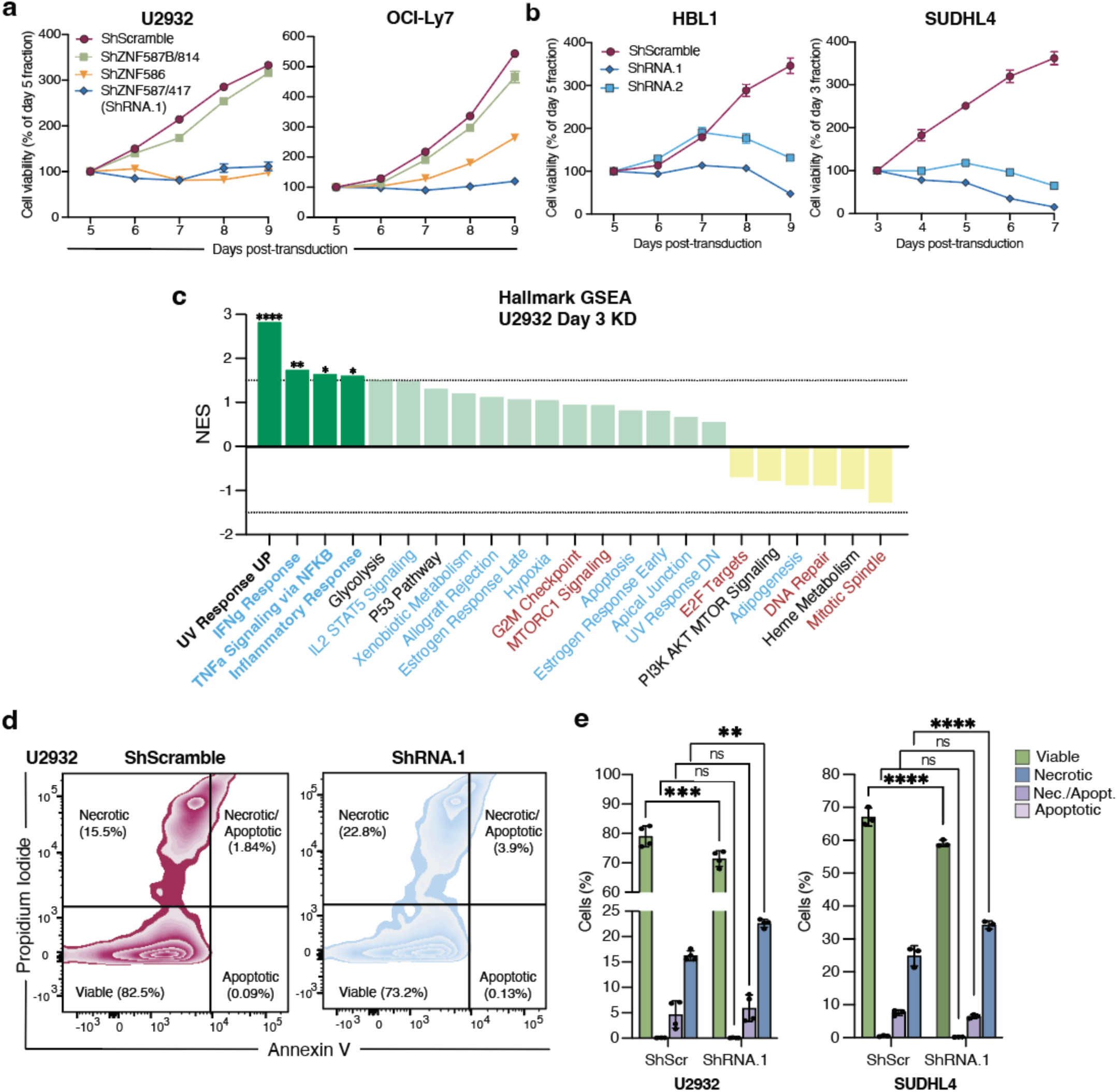
ZNF587/417 depletion impairs cell growth and viability. (a) MTT proliferation assays of U2932 and OCI-Ly7 cells upon LV transduction with shRNAs targeting *ZNF587B/814*, *ZNF587/417*, and *ZNF586* or a control shRNA (shScr). (b) MTT proliferation assays for HBL1 and SUDHL4 upon LV transduction with 2 different shRNAs targeting *ZNF587/417* (shRNA.1/.2) or a control shRNA. (c) Waterfall plot of Hallmark GSEA signatures from RNA-seq data ranked by their Normalized Enrichment Score (NES)^93^. Enriched signatures (NES>0) are highlighted in green and depleted signatures (NES<0) are in yellow. The p-value cutoff <0.05 is indicated by a dotted line. Hallmark gene sets enriched in KZFP^High^ or KZFP^Low^ DLBCLs are highlighted in dark red and blue, respectively. (d) Representative flow cytometry plot using Annexin V-APC/PI staining for cell death characterization in U2932 cells. (e) Bar plots showing the percentage of cells undergoing early (PI-/aV+) *vs*. late apoptosis (PI+/aV+) and necrosis (PI+/aV-) in U2932 and SUDHL4 cells, 3 days post-transduction with shRNA.1 targeting *ZNF587/417* or a control shRNA (mean ± SD of independent duplicates). Statistics: Two-Way ANOVA with Sidak’s multiple comparison test.

In order to understand the mechanisms underlying this observation, we performed RNA sequencing (RNA-seq) on U2932 cells harvested 3 days after *ZNF587/417* KD. We identified 332 differentially expressed genes (DEGs, fold change > 2 and FDR <0.05), 178 (53.6%) of which were downregulated (Fig. 3c). Gene set enrichment analysis (GSEA) revealed UV response UP as the most affected hallmark amongst DEGs. It encompassed several cyclins (*CCND3, CCNE1*), cyclin-dependent kinase inhibitors (*CDKN1A, CDKN1B, CDKN1C*), and TP53-responsive genes (*TP53I3*) (Fig. S2e). Reminiscent of the form of cell death induced by irradiation^72^, Annexin V and Propidium Iodide (PI) staining revealed an increase in PI+/AV-cells, a pattern indicative of necrosis, upon *ZNF587/417* KD in U2932 and most prominently SUDHL4 cells (Fig. 3f). Thus, ZNF587/417 depletion impaired proliferation and triggered necrotic cell death across DLBCL cell lines.

### ZNF587/417 depletion leads to a broad downregulation of KZFPs and alters the heterochromatin landscape of lymphoma cells

ZNF587 and ZNF417 are both KAP1-binding KZFPs that secondarily recruit the methyltransferase SETDB1 responsible for catalyzing the deposition of H3K9me3, a repressive histone mark promoting heterochromatin formation at targeted loci^71^. The two paralogs recognize, almost exclusively, overlapping sets of primate-restricted TEs, including members of the ERVK and ERV1 classes of ERVs as well as many SVA integrants^31,71^. To ask if the phenotype observed upon *ZNF587/417* KD cells correlated with changes in their chromatin status, we performed genome-wide profiling of H3K9me3 using CUT&Tag^73^ in U2932 cells at day 3 of KD. Similar peak numbers (n=26’841) and locations were detected in the control and KD cells, but in ZNF587/417-depleted cells peak intensity significantly decreased at 1’577 and increased at 755 locations, most of the latter in pericentromeric regions (89.9% vs. 10.07%, p <0.0001; Fig. 4a). Measurement of H3K9me3 across all called peaks confirmed a marked increase at pericentromeric regions and telomeres, contrasting with a global decrease across the rest of the genome following *ZNF587/417* KD (Fig. 4b).

**Fig. 4:**
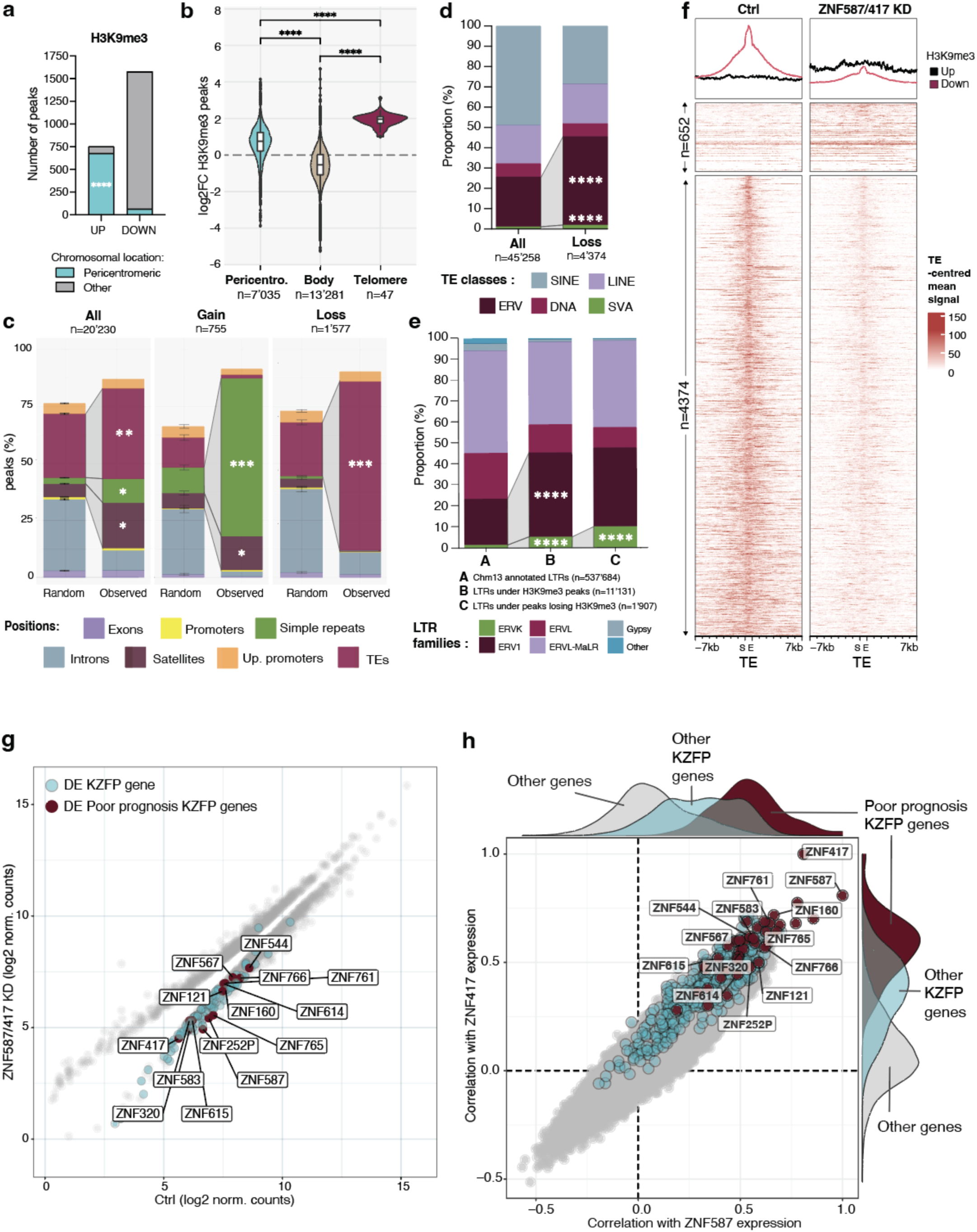
ZNF587/417 depletion alters the heterochromatin landscape. (a) Number of Cut&Tag peaks with significant (FDR <0.05) gain (UP) or loss (DOWN) of H3K9me3, 3 days post-transduction with shRNA.1 targeting *ZNF587/417* or a control shRNA (two independent duplicates). (b) Violin plots of the distribution of the log2 fold change (shRNA.1 vs. control) of H3K9me3 signal over called peaks in the listed genomic features. Statistics: Two-sided Mann–Whitney U-test. (c) Observed genomic distribution of all H3K9me3 peaks (left) and peaks displaying a significant gain (center) or loss (right) of H3K9me3 signal vs. randomized peaks, over listed genomic features (mean value of dupicates +/− SD from 1000 permutations; FDR <0.05). Genomic positions overlapping with >50% an H3K9me3 peak were assigned in the following order: gene promoters (up to 1000bp upstream from the transcription start site, TSS), up.promoters (up to 5000bp upstream from TSS), exons, TEs, satellites, simple repeats, and introns. Statistics: Chi-squared test, alternative: greater (d) Proportion of TE classes overlapping all H3K9me3 called peaks (left) or peaks with a significant loss of H3K9me3 (FDR <0.05, right). Asterisks indicate a significant enrichment compared to all H3K9me3 called peaks. Statistics: Two-tailed Fisher’s Exact Test. (e) Same as (d) over LTR families, including peaks annotated in the T2T-CHM13v2.0 genome release (left). Statistics: Two-tailed Fisher’s Exact Test (f) Profile heatmaps centered on TEs (+/− 7kb) displaying significant change in H3K9me3 Cut&Tag signal (>50% overlap; mean signal coverage from duplicates). (g) Scatter plot of RNA-seq data from ZNF587/417 KD versus control cells at day 3 of KD in U2932, outlining DEGs (grey dots, FDR <0.05), poor-prognosis KZFP genes (dark red dots) and other KZFP genes (turquoise dots). (h) Scatter plot of Spearman’s correlation coefficients between ZNF417 (y-axis) or ZNF587 (x-axis) expression, and each gene expressed amongst the 633 DLBCLs analyzed transcriptomes (n= 23’560), outlining poor-prognosis KZFP genes (dark red dots), other KZFP genes (turquoise dots), and other genes (grey dots), with density plots showing their distribution on each side.

While H3K9me3 peaks in control U2932 cells were enriched at TEs, simple repeats and satellite repeats, the KD triggered a preferential loss of this repressive mark at TEs and a gain at the other two categories of repeats (Fig. 4c). Amongst TEs, the most significant losers H3K9me3 upon *ZNF587/417* KD belonged to the ERV and SVA subgroups (Fig. 4d), notably ERVK integrants previously identified as the preferential ERV targets of these KZFPs ^71^ (Fig. 4e). When some increase in H3K9me3 was noted over TE inserts, it likely reflected spreading of this mark from surrounding satellite/simple repeats (Fig. 4c/f). Accordingly, peaks gaining H3K9me3 were on average substantially wider than TE-centered peaks losing this mark (Fig. S3a). Still, the loss of H3K9me3 over a range of TEs extending beyond the sole known targets of ZNF587/417 suggested that additional factors contributed to this process. Confirming this suspicion, our RNA-seq analysis revealed the downregulation of another 100 *KZFP* genes (FDR <0.05) in U2932 cells 3 days after ZNF587/417 KD (Fig. 4g), including 12 poor-prognosis *KZFP* gene hits. By contrast, only one *KZFP* gene was found upregulated in *ZNF587/417* KD cells at this time point. This downregulation was even broader at day 6 of KD, encompassing 184 and 133 *KZFP* genes (FDR <0.05) besides *ZNF587* and *ZNF417* in U2932 and OCI-Ly7 cells, respectively (Fig. S3b). While the reason for such broad downregulation is unknown, we observed that the expression of *ZNF587/417* was highly correlated with that of other *KZFP* genes in our DLBCL cohort (Fig. 4h), pointing to common or cross-regulatory mechanisms within this gene family.

### ZNF587/417 depletion triggers replicative stress in lymphoma cells

Our data pointed to a link between ZNF587/417 depletion and a UV irradiation-like transcriptional response. To probe this issue further, we measured the impact of *ZNF587/417* KD on Ser-139 H2AX phosphorylation (*γ*H2AX) in U2932 and OCI-Ly7 cells 3 days after KD, which revealed a marked increase in this mark of DNA damage response (DDR) (Fig. 5a/S5a). In order to determine whether this DDR was concentrated in particular regions of the genome, we performed CUT&Tag for *γ*H2AX in U2932 cells after 3 days of *ZNF587/417* KD. Due to the known critical role of peri-/centromeric regions and telomeres in chromosomal integrity, we used the new Telomere-to-Telomere assembly of the *CHM13* cell line (T2T-CHM13)^74^ for our analysis. Peak numbers (n=7’635) and locations were similar in control and KD cells, but *γ*H2AX levels were significantly increased over 311 loci and decreased over 101 loci upon *ZNF587/417* KD (Fig. 5b), including 14 upregulated and 5 downregulated peaks located at pericentromeric regions (Fig. 5c). Although the number of peaks with a significant *γ*H2AX increase was not enriched at those regions, the peaks located at pericentromeric regions (n=480) and telomeres (n=42) presented a higher global fold increase in *γ*H2AX compared to other peaks upon *KZFP* KD (Fig. 5d). This reflected a significant enrichment in *γ*H2AX-gaining peaks overlapping with simple repeats compared with other *γ*H2AX-bearing loci (Fig. 5e). These results suggested that, when *ZNF587/417* are depleted, DNA damage was exacerbated at preexisting *γ*H2AX hotspots rather than at TE loci targeted by these KZFPs. Since exacerbated DDR at preexisting *γ*H2AX hotspots has been described with RS inducers such as hydroxyurea or aphidicolin^75^, we characterized DNA replication fork progression at the genome-wide level using <u>Tr</u>ansferase-<u>A</u>ctivated <u>E</u>nd <u>L</u>igation <u>seq</u>uencing (TrAEL-seq). We observed that *ZNF587/417* KD cells exhibited a global loss of replication polarity, indicating a lower frequency of diverging replication forks related to an increase in the number of replication origins (Fig. 5f). The excessive firing of replication origins is a known source of RS, notably due to the exhaustion of limiting factors necessary for replication fork progression^76,77^ and collisions between replication and transcription^78^, but it can also be a protective mechanism from RS triggered by primary stalling of replication forks^79^. In support of the former hypothesis, we observed a strong correlation between H3K9me3 changes and replication fork polarity in *ZNF587/417* KD cells (Fig. 5g), arguing in favor of a link between these two alterations. Strikingly, genomic regions with a loss in H3K9me3 exhibited a decrease in replication directionality, whereas the opposite was observed where H3K9me3 was gained (Fig. 5h), that is, predominantly at pericentromeric regions. To probe if the cell growth impairment observed upon ZNF587/417 depletion was due to RS, we assessed DNA replication by measuring the incorporation of the thymidine analog 5-Ethynyl-2′-deoxyuridine (EdU) in control and knockdown U2932 and SUDHL4 cells by flow cytometry. It revealed a decrease in EdU incorporation, along with a proportional increase of non-replicating cells during the S phase of the cell cycle (Fig. 5i), thus confirming RS-related arrest.

**Fig. 5:**
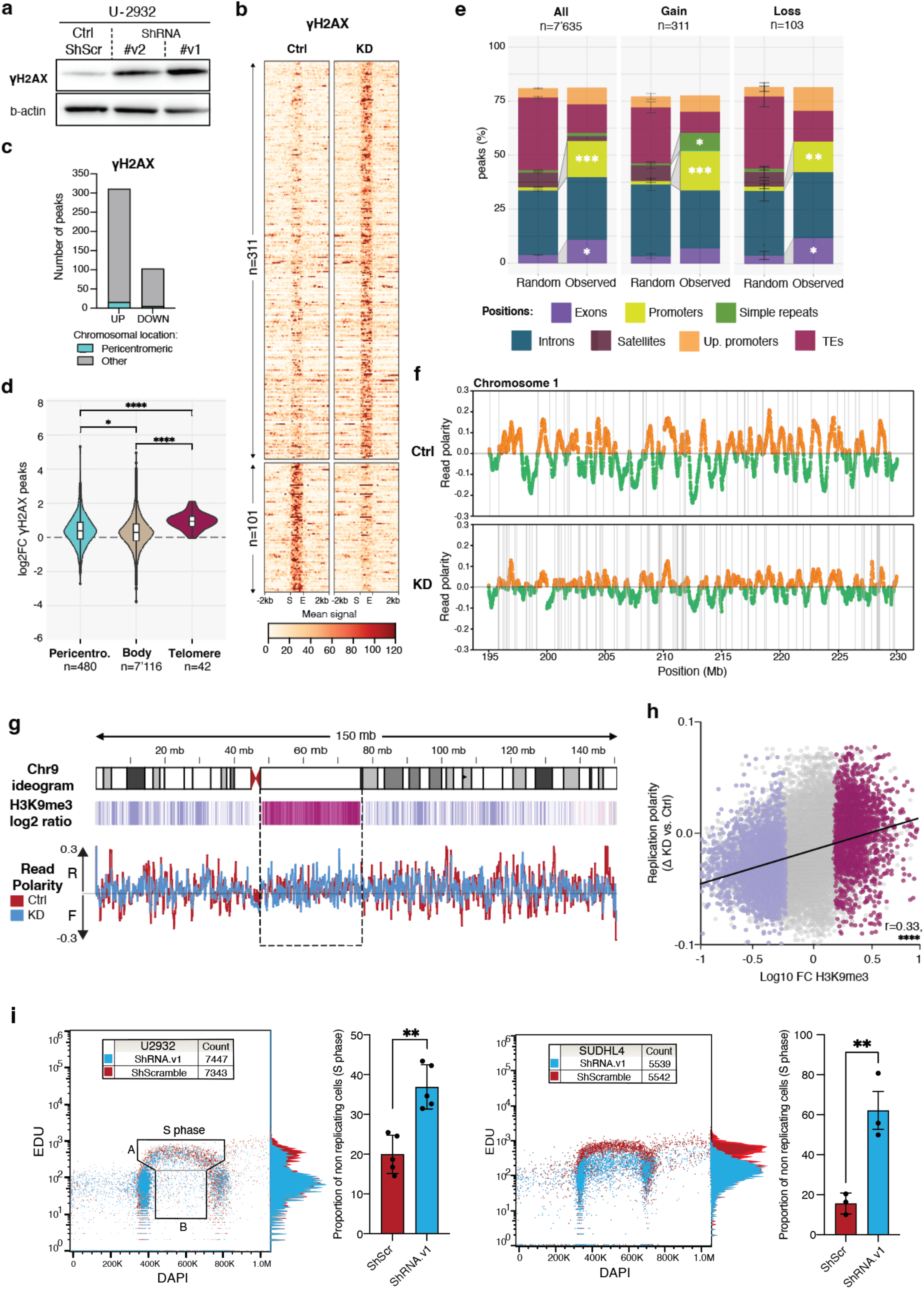
ZNF587/417 depletion triggers replicative stress. (a) Western blot analysis of *γ*H2AX in U2932 cells 3 days after transduction with 2 different shRNAs targeting *ZNF587/417* (shRNA.v1/.v2) or a control shRNA (shScr). *β*-actin was used as a loading control. (b) Profile heatmaps of significantly changing *γ*H2AX Cut&Tag peaks (p-val. <0.05) in U2932 control and KD conditions, 3 days post-transduction (mean signal coverage from duplicates). (c) Number peaks with a significant gain (UP) or loss (DOWN) in *γ*H2AX in U2932 KD cells vs. control (p val <0.05). (d) Violin plots of the distribution of the log2 fold change (shRNA.1 vs. control) of *γ*H2AX signal over called peaks in the listed genomic features. Statistics: Two-sided Mann–Whitney U-test. (e) Observed genomic distribution of all *γ*H2AX peaks (left) and peaks displaying a significant gain (center) or loss (right) of *γ*H2AX signal, vs. randomized peaks, over listed genomic features (mean value of duplicates +/− SD from 1000 permutations; FDR <0.05). Genomic positions overlapping with >50% a *γ*H2AX peak were assigned in the following order: gene promoters (up to 1000bp upstream from the transcription start site, TSS), up.promoters (up to 5000bp upstream from TSS), exons, TEs, satellites, simple repeats, and introns. Statistics: Chi-squared test, alternative: greater (f) TrAEL-seq read polarity plots over the chromosome 1 in U2932 control and KD conditions, 3 days post-transduction. Read polarity was calculated using a 250 kb sliding window spaced every 10 kb across the genome – *see methods.* Orange and green points represent sliding windows with a reverse or forward strand bias, respectively. Vertical grey lines represent shifts from forward to reverse bias, highlighting strong replication origins. (g) Top: Heatmap track of log2 ratio of Cut&Tag H3K9me3 signal between control and KD conditions surrounding the pericentromeric region of the chromosome 9. Significant H3K9me3 gain is highlighted by the dashed rectangle. Bottom: Overlapped read polarity plots of control (dark red) and KD (blue) conditions. (h) Scatter plot of replication polarity *vs.* H3K9me3 Cut&Tag signal changes. X-axis: log10 fold change of all H3K9me3 peaks called in U2932 cells. Y-axis: Replication polarity difference between KD and control conditions calculated from the absolute value of the percentage of reverse reads in a 2 Mb window surrounding the center of each H3K9me3 peak. Peaks with a significant gain in H3K9me3 are shown in dark red, the one with a significant H3K9me loss in turquoise blue, and others in grey. (i) Representative flow cytometry analysis of the replication signal (EdU incorporation intensity, y-axis) and DNA content (DAPI staining, x-axis) in U2932 and SUDHL4 KD (blue) *vs.* control cells (dark red), 3 days post-transduction. Y-axis side histogram shows changes in EdU intensity between KD (blue) and control (dark red) conditions. S phase cells are defined as events located within the A+B gates; A area= replicating cells, B area = non-replicating cells. Right: Barplots depicting the mean proportion (+/−SD) of non-replicating S phase cells calculated as follows: (B/(A+B)) *100 from 5 and 3 biological replicates in U2932 and SUDHL4, respectively. Statistics: Student’s t-test.

### ZNF587/417 depletion leads to cell-intrinsic inflammation

At day 6 of KD, 732 genes were upregulated and 915 downregulated (FC>2, FDR <0.05; Fig. 6a), including 195 the 332 (58.7%) genes already noted as differentially expressed at day 3 (Fig. 6b). The transcriptome was then characterized by the upregulation of IFN-responsive and other inflammatory genes, and by an enrichment in Hallmark gene sets previously noted in KZFP^Low^ DLBCL samples (Fig. S5a). Interestingly, IFN-related DNA damage signature (IRDS) genes, such as *STAT1*, *IRF7*, *IFIT1/3*, *MX1*, *ISG15*, *IFI44* and *OASL* family members were upregulated at day 6 and 10, suggesting that the observed inflammatory response was at least partially related to genomic instability (Fig. 6c). TEs being well known sources of cell-intrinsic inflammation, we monitored their transcription during these events. Even though some TE loci were detected as mildly expressed at day 6, frank TE upregulation became apparent only at day 10 post-KD (Fig. S5b). Furthermore, the TEs significantly upregulated at day 6 belonged to the SINE group, probably induced by the ongoing genotoxic stress^80^. In contrast, most TEs significantly induced at day 10 were ERVs (Fig. S5c), notably HERV9, HERV9N, HERV9NC, and LTR12C subfamily members of the primate-specific ERV1 family harboring ZNF587/417-binding motifs (Fig. S5d). Of note, these ERV subsets already displayed an increased expression at day 6, albeit not reaching statistical significance at the TE family level (Fig. S5c). Irrespectively, our transcriptome analyses revealed that IFN/inflammatory-related genes were already upregulated at day 3 of *ZNF587/417* knockdown, that is, still in the complete absence of TE induction (Fig. 3c). Therefore, TE transcripts and their products could not be the source of the inflammation initially observed upon KZFP depletion.

**Fig. 6:**
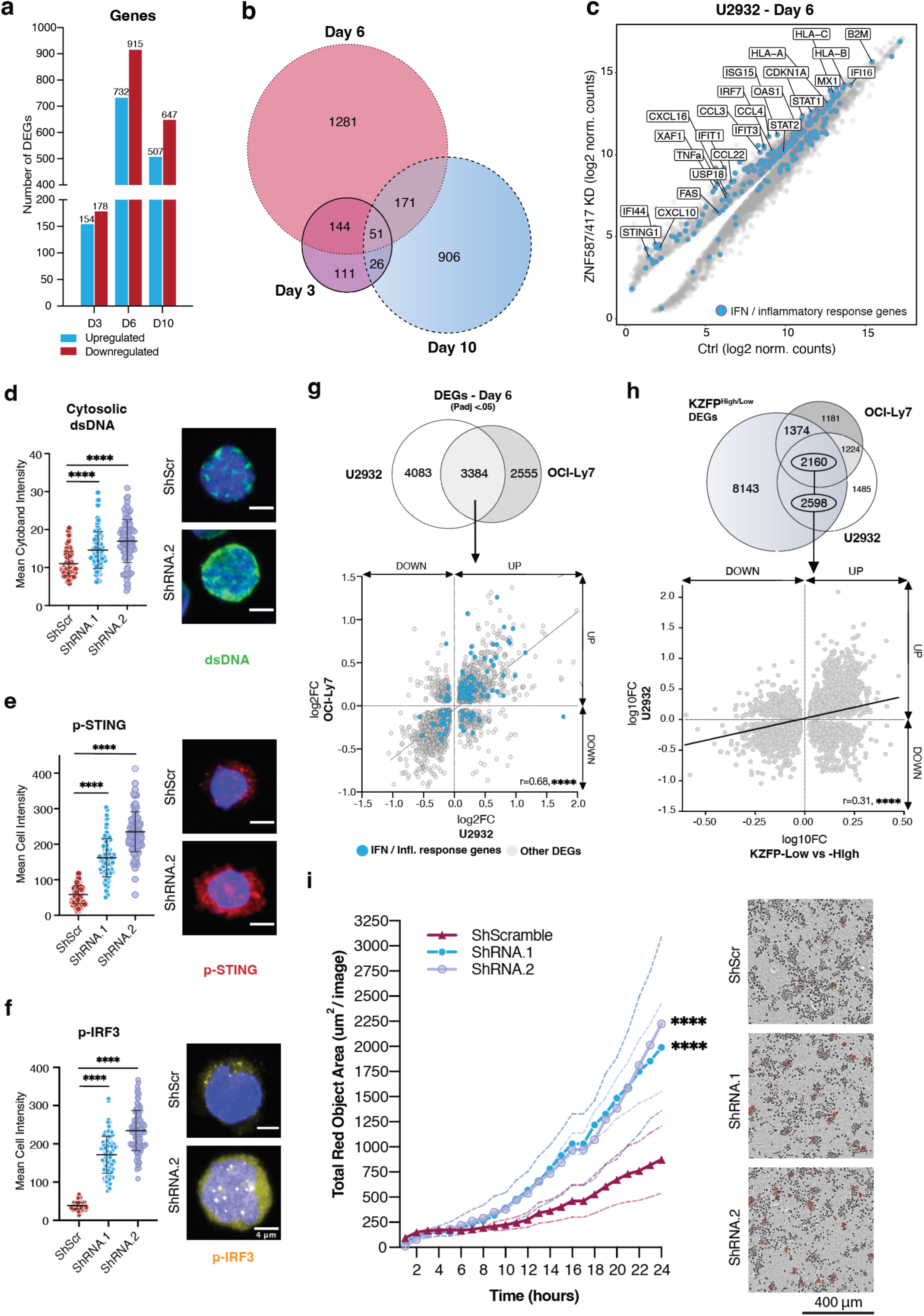
ZNF587/417 depletion leads to cell-intrinsic inflammation. (a) Number of DEGs of RNA-seq analysis performed at day 3 (n=2), day 6 (n=3), and day 10 (n=2) of ZNF587/417 KD vs control U2932 cells. (b) Euler diagrams of the overlap of DEGs upon ZNF587/417 KD in U2932 cells at each time point (day 3, 6, and 10). (c) Scatter plot of log2 normalized counts of ZNF417/587 KD cells vs. control U2932 at day 6, outlining DEGs (grey dots) among which genes belonging to type I/II IFN and Inflammatory response Hallmark gene sets, Interferon Signaling Reactome, and cellular response to type I, II and III IFN gene ontology terms are depicted in blue. (d) Representative images and dot plot showing the mean intensity of cytosolic double stranded DNA per cell, measured on z-stack immunofluorescence images (n *≥*80 cells per condition). Statistics: Two-sided Mann–Whitney U-test. (e,f) same as (d) for phosphorylated STING (e) and phosphorylated IRF3 (f) signal respectively. (g) Top: Venn diagrams showing the overlap of DEGs upon ZNF587/417 KD in U2932 and OCI-Ly7 at day 6. Bottom: Scatterplot of Log2 fold changes of DEGs shared between U2932 (x-axis) and OCI-Ly7 (y-axis) KD cells. Blue dots highlight the genes belonging to IFN-/Inflammatory response terms detailed in (c). The best-fit line in grey results from the linear regression analysis of U2932 log2 fold changes onto OCI-Ly7 log2 fold changes. (h) Top: Venn diagrams showing the overlap of DEGs upon ZNF587/417 KD in U2932 and OCI-Ly7 cells at day 6 and genes discriminating KZFP^High^ and KZFP^Low^ DLBCLs. Bottom: Scatterplot of log2 fold changes of DEGs shared between genes discriminating KZFP^High^/KZFP^Low^ DLBCLs (x-axis) and U2932 KD cells (y-axis). The best-fit line in black results from the linear regression analysis of KZFP^High^ vs KZFP^Low^ log2 fold changes onto U2932 KD log2 fold changes. For this panel, DEGs were defined with a FDR <0.05 and a fold change >2. r: correlation coefficient. (i) Phagocytosis assay of U2932 pHrodo-labeled cells co-cultured with M1-polarized macrophages. Representative 24h course of pHrodo signal quantification using InCucyte time-lapse imaging of ZNF587/417 KD cells with two different shRNAs (shRNA.1 in blue and shRNA.2 in turquoise blue) and control cells (shScramble in dark red). Total pHrodo cell area per image acquired was calculated for each time-point and plotted as a time course for each condition. Right: representative images of respective conditions. Statistics: Two-way ANOVA followed by Bonferroni correction. Asterisks indicate significant differences at 24h time-point compared to controls.

The accumulation of cytosolic DNA molecules due to endogenous or exogenous genotoxic stress is another known trigger of an IFN response, occurring via activation of the cGAS-STING DNA sensing pathway^81–83^ by micronuclei or free DNA fragments generated respectively in the context of lagging chromosomes^84^ or replication defects^83,85^. At day 3 of *KZFP* KD, U2932 cells presented a significant accumulation of cytosolic dsDNA (Fig. 6d) as well as of the dsDNA sensing intermediate mediator phospho-STING (Fig. 6e), together with the phosphorylation and nuclear translocation of IRF3, one of its main downstream effectors (Fig. 6f). Accordingly, upregulation or IRF3 target genes was recorded at day 6 of *ZNF87/417* KD (Fig. S5e). Furthermore, transcripts encoding CXCL10, a chemokine promoting the attraction of mononuclear phagocytes, T cells, and Natural Killer (NK) cells, were upregulated starting from day 3 up to day 10 following *ZNF587/417* KD, and we could confirm increased levels of this cytokine in the cell supernatant (Fig. S5f). Induction of a cell-intrinsic inflammatory response was not restricted to ABC cells, as upregulation of IFN-related responsive genes similarly occurred in GCB-related OCI-Ly7 cells following *ZNF587/417* KD (Fig. 6g/S5g), thus further highlighting that the effect of KZFP depletion was similar across different cellular contexts. Furthermore, upon intersecting genes differentially expressed at day 6 post-KD in U2932 or OCI-Ly7 cells with the genes defining the KZFP^High^ and KZFP^Low^ groups of the DLBCL cohort, we found that *ZNF587/417* KD induced a shift of the cell lines towards a KZFP^Low^-like transcriptome, supporting an important role for ZNF417 and ZNF587 in the gene expression profile of poor-prognosis DLBCL (Fig. 6h). In order to assess if these changes would lead to an increased susceptibility to innate immunity clearance, we tested the phagocytosis of human M1 polarized macrophages exposed to ZNF587/417 KD cells during 24 hours between day 4 and 5 of KD (Fig. 6i). Indeed, M1 macrophages exhibited a stronger phagocytic ability against ZNF587/417 KD cells compared to control cells, suggesting that in the clinics ZNF587/417 blockade could foster the phagocytic clearance DLBCL residual disease. Interestingly, the transcriptome of the U2932 KD cells started reverting towards a KZFP^High^ profile at day 10, despite a persistent *KZFP* downregulation, with re-induction of cell cycle-related genes such as E2F targets and G2M checkpoint genes, and downregulation of transcripts related to EMT, TNF*α* signaling via NF-kB, and hypoxia (Fig. S5h). This suggested ongoing transcriptional reprogramming, probably driven by inflammation and DNA repair, with progressive establishment of a resistant phenotype.

### ZNF587/417 depletion modifies the antigenic profile of lymphoma cells

Our data indicate that the RS induced by *ZNF587/417* KD triggers an inflammatory response through the cytosolic release of DNA fragments and their sensing by the cGAS-STING pathway. This type of response normally leads to increased expression of *HLA-I* genes, which play a pivotal role in the immunogenicity of cancer cells. Accordingly, in U2932 cells, *HLA-C* gene expression started to be significantly upregulated at day 3 of *ZNF587/417* KD (Fig. S2e), followed by *HLA-A*, *HLA-B*, and beta-2-microglobulin (*B2M*) at day 6 (Fig. 6c), while in OCI-Ly7 *HLA-B* and *HLA-C* transcripts were augmented only at this later time point (Fig. S5g). Corroborating these transcriptomic changes, U9932 cells displayed a 2-fold increase in surface expression of HLA-A, -B, and -C at KD day 10, even though at that point their source transcripts were no longer upregulated (Fig. 7a). We went on to assess whether the upregulation of HLA-I observed in *ZNF587/417* KD cells was associated with an alteration of their HLA-bound peptidome. For this, HLA-bound peptides of KD day 10 and control U2932 cells were eluted and sequenced by mass spectrometry. We found 1’367 peptides derived from 1050 different proteins (∼14.3% of the detected immunopeptidome and 23% of presented proteins) to be significantly more and 1154 peptides from 876 proteins (∼12% of the detected immunopeptidome and 19.2% of presented proteins) to be less abundant in *ZNF587/417* KD cells (*P* <0.05; Fig. 7b). In accordance with the transcriptome data, most of the peptides enriched in *ZNF587/417* KD cells corresponded to inflammatory gene products, while KZFP-derived peptides were prevalent amongst depleted entities (Fig. 7b). Furthermore, the peptide repertoire of U2932 KD cells was altered, with noticeable changes in peptide length: 8 and 9-mers made up 90.5% of the peptides enriched in KD cells but only 65.9% of the peptides enriched in control cells, and conversely 10- and 11-mers constituted only 8.8% of the peptides enriched in U2932 KD *versus* 29.4% in control cells (Fig. 7c). We hypothesized these changes to be linked to differences in HLA allele-specific expression. By using previously described HLA binding prediction methods (Fig. 7d), we found that the percentage of peptides predicted to bind HLA-B (69.4% *vs* 41.8%) and -C (7.7% *vs* 3.5%) was increased in U2932 KD cells compared to control cells, where peptides were predicted to bind mostly HLA-A03:01 (50.1%), an allele recently associated to poor responses to immune-checkpoint blockade^86^. The *ZNF587/417* KD increased also the proportion of hydrophobic (leucine) while decreasing that of amphipathic (lysine) residues at the C-terminus of the presented peptides (Fig. 7c), suggesting underlying alterations of the proteasome degradation pathway and changes in HLA binding affinity. Therefore, we compared the immunogenicity of the eluted peptides using the Immune Epitope Database (IEDB) analytic tool^87^, which predicted that immunopeptides eluted from U2932 KD cells were on average more immunogenic than that of control cells (Fig. 7e). Together these results indicate that *ZNF587/417* KD reshuffles the immunopeptidome of lymphoma cells resulting in increased immunogenicity.

**Fig. 7:**
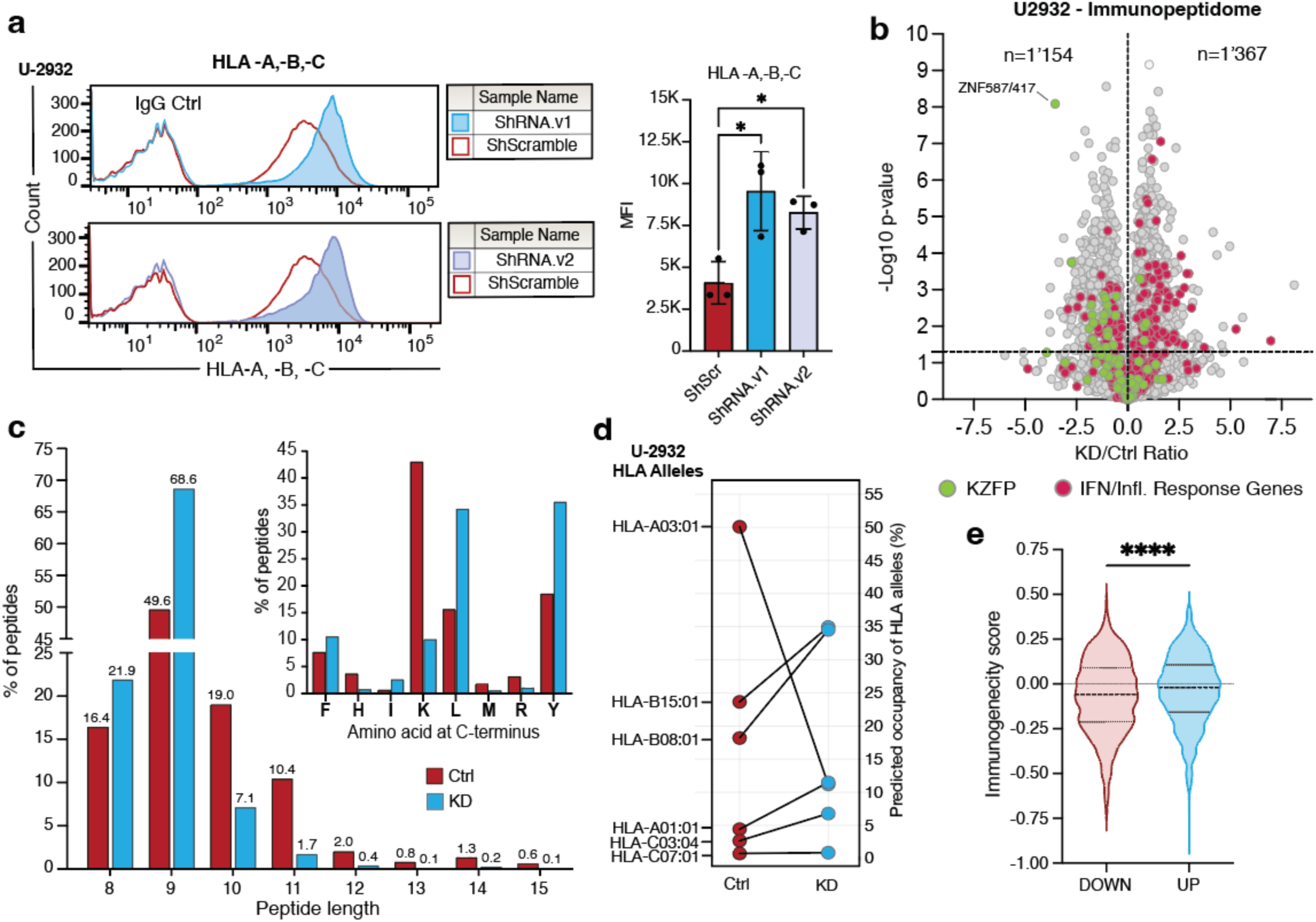
ZNF587/417 depletion modifies the antigenic profile of lymphoma cells. (a) Representative flow cytometry histograms (left) and bar plot quantification (right) of median fluorescence intensity (MFI) of HLA class I (HLA-A, -B, -C) expression on U2932 KD and control cells after 10 days with two different shRNAs targeting *ZNF587/417* (shRNA.1/.2) or a control shRNA (shScr). Statistics: Student’s t-test (mean of MFI +/− SD of independent triplicates). (b) Volcano plot showing changes in the abundance of HLA-I−bound peptides between ZNF587/417 KD (shRNA.1/.2) and control cells from 3 independent experiments. The horizontal dashed line indicates a p-value threshold of 0.05. n, number of enriched and depleted peptides. Peptides derived from gene products belonging to type I/II IFN and Inflammatory response Hallmark gene sets, Interferon Signaling Reactome, and cellular response to type I, II, and III IFN gene ontology terms are highlighted in dark red. Peptides derived from KZFPs are highlighted in green. Statistics: Student’s t-test. (c) Bottom left: Differences in the proportion of peptides of different lengths between control (dark red) and KD cells (blue). Top right: Bar plot showing the differences in the percentage of peptides with listed C-terminal amino acid between control (dark red) and KD cells (blue). (d) Line plot showing the changes in the proportion of predicted occupancy of HLA-alleles present in U2932 cells between control (dark red) and KD cells (blue). (e) Violin plots showing the predicted immunogenicity scores of eluted peptide ligands enriched in control (DOWN, red) and KD cells (UP, blue) from the IEDB analytic tool.

## DISCUSSION

Genomic instability confers cancer cells with evolutionary advantages by generating clonal diversity, which facilitates escape from immune surveillance and chemotherapeutic agents. Yet it has to be balanced in order to avoid mitotic catastrophes and anti-tumoral immune responses. Constitutive heterochromatin is a dynamic state reestablished after each cell division through epigenetic memory^88–90^. The present work demonstrates that KZFPs, the TE-targeting partners of the KAP1-SETDB1 complex, play an important role in the maintenance of heterochromatin in cancer cells, protecting these from excessive RS, secondary inflammatory responses and other immunologically detectable alterations. Our results further identify ZNF587 and ZNF417, two primate-specific paralogs, as leading pro-oncogenic KZFPs, and their upregulation as a strong negative prognosis indicator in DLBCL.

Recent studies have revealed that chromosomal instability (CIN) is a main driver of cancer progression and metastasis, and pointed to the induction of the cGAS-STING dsDNA sensing pathway as an important mediator of this process^11^. CIN^High^ tumors are typically described as carrying a poor prognosis and displaying an activation of genes related to IFN/inflammatory responses, EMT and TNF*α* signaling^11^. Here, we found KZFP^High^ DLBCLs to exhibit a higher average number of CNAs than their KZFP^Low^ counterparts, yet their transcriptome to be characterized by an upregulation of DNA repair genes but CIN^Low^ expression signatures linked to cell cycle, Myc target, and OxPhos-related genes. Thus, KZFP^High^ DLBCL cells presumably combine higher degrees of genetic instability with an increased ability both to attenuate RS and to dampen inflammatory responses. As a consequence, the stabilization of their chromatin via upregulation of critical controllers of repetitive DNA results both in increasing the efficiency of their clonal expansion and diversification and in facilitating their immune evasion. ZNF587/417 recognize evolutionarily recent ERVs and SVAs, capable of robust transcriptional influences notably manifested during brain development^31,71^. In most tissues, *cis*-acting regulatory sequences embedded in these retroelements are maintained in a repressed state through a mix of histone and DNA methylation, the latter perpetuated across mitosis through the action of maintenance DNA methyltransferases^30^, without the need for sequence-specific repressors. Our data suggest a model whereby, perhaps due to the state of DNA hypomethylation characteristic of cancer cells, maintenance of heterochromatin at these TE loci becomes strongly dependent on high levels of histone methylation-inducing KZFPs recognizing their sequence in order to prevent inflammatory responses.

The importance of KZFPs in DLBCL pathogenesis is supported by the recurrence of amplifications of the distal portion of Chr19q in this lymphoid malignancy^91,92^. This phenomenon had been so far attributed to the presence in this chromosomal segment of *SPIB*, a gene encoding a repressor of the IFN*β* gene activator IRF7^91^. However, SPIB acts in a COO-restricted manner^93^, whereas Chr19q amplifications have been detected in both ABC and GCB DLBCLs and demonstrated to correlate with poor prognosis independently from the lymphoma COO in prospective cohorts^92^. Our results thus strongly suggest that this chromosomal amplification is driven by the pro-oncogenic potential of *ZNF587/417* and other components of *KZFP* gene cluster 11, as well as perhaps by the additional presence in the sub-telomeric region of Chr19q of *KAP1*, which not only is an essential cofactor of these KZFPs but also itself a negative prognostic factor in non-Hodgkin B cell lymphoma^94^.

Our finding that *γ*H2AX accumulates at hotspots of endogenous DNA damage upon *ZNF587/417* KD is reminiscent of chromatin alterations detected in lymphoblastoid cell lines exposed to chemical inducers of RS^75^. Moreover, the reshuffling of the immunopeptidome observed in lymphoma cells surviving the acute genotoxic stress triggered by *ZNF587/417* KD closely resembles changes observed upon exposure of tumor cells to non-cytotoxic doses of the nucleoside analogue gemcitabine^95^.

Together, these data indicate that methods aimed at inactivating ZNF587/417 could mimic the action of chemotherapeutic agents. Our finding that normal cells do not undergo genotoxic stress when depleted in these KZFPs suggests that approaches targeting these TE controllers might have a very favorable therapeutic index, further warranting efforts geared toward their development.

## MATERIAL AND METHODS

### Cell culture

OCI-Ly7 and U2932 human lymphoma cell lines were obtained from DSMZ – German Collection of Microorganisms and Cell Cultures, Braunschweig, Germany (www.dsmz.de). The other DLBCL cell lines, namely SUDHL4 and HBL1 were kindly supplied by the laboratory of Professor Elisa Oricchio (EPFL, Swiss Institute for Experimental Cancer Research) and typed using short tandem repeat (STR) profiling by Microsynth cell line authentication service. U2932, HBL1, and SUDHL4 cells were grown in RPMI 1640 (Gibco) with 10% fetal bovine serum and 1% penicillin-streptomycin. OCI-Ly7 cells were grown in IMDM (Gibco) with 20% fetal bovine serum (FBS) and 1% penicillin-streptomycin. All cell lines were grown at 37°C and 5% CO2. HEK293T cells were grown in DMEM (Gibco) supplemented with 10% FBS, 1% penicillin-streptomycin, and 2mM of L-glutamine. Primary normal dermal fibroblasts were cultured in Fibroblast Basal Medium (ATCC) supplemented with Fibroblast Growth Kit-Low Serum (ATCC). Human M1-polarized macrophages and M-CSF were purchased from StemCell. After thawing, M1 macrophages were resuspended in complete RPMI 1640 (Gibco) medium with M-CSF (StemCell) at a final concentration of 50 ng/ml.

### KZFP knockdown experiments

The short hairpin sequences targeting ZNF587/417 were designed as described^71^. ZNF587/417 shRNAs are targeting the following sequences: 5’-GCAGCATATTGGAGAGAAATT-3’ (shRNA.1) and 5’-AGTCGAAAGAGCAGCCTTATT-3’ (shRNA.2). The other shRNAs were purchased from Sigma-Aldrich, with shRNAs against ZNF587B/814 and ZNF586 targeting the following sequences: 5’- CCTTCTAAGCAGAGTATTTAT- 3’ (shZNF587/814), 5’- GCTTATACATCTAGTCTCATT-3’ (shZNF586). An shRNA with the non-targeting sequence 5’-CAACAAGATGAAGAGCACCAAG-3’ (shScramble, shScr) was used as a negative control in all experiments. Lentivectors (LVs) were produced as described on the following website: http://tronolab.epfl.ch. Lymphoma cell lines and primary fibroblasts were transduced with a MOI of 10. 48h post-transduction, cells were selected for 3 days using 3 μg/mL of puromycin. For experiments performed between day 2 to 4, cells were used without selection while doubling the multiplicity of infection (MOI of 20).

### Cell proliferation assays

The tetrazolium bromide (MTT) assay was used to determine cell proliferation for 4 days. Cells were plated after 5 days of LV transduction and 3 days of puromycin selection, except for SUDHL4 cells which were plated 3 days after LV transduction. Briefly, 100μL of cells were seeded in 96-well plates at a concentration of 5k/well for primary dermal fibroblasts, 20k/well for HBL1, and 40k/well for U2932, OCI-Ly7, and SUDHL4 cells. On each consecutive day, cells were incubated with 5 mg/ml MTT for 3h at 37°C. The reaction was stopped by the addition of 100μL of 10% Ultrapure SDS (Invitrogen) to dissolve precipitated crystals. After overnight incubation at 37°C, the absorbance was measured on a microplate spectrophotometer at 570 nm while subtracting the background absorbance measured at 690 nm.

### Annexin V/propidium iodide cell death detection assays

Cell death was determined using flow cytometry through the quantification of cell surface annexin V-APC (Biolegend) and propidium iodide (PI, Biolegend) co-staining. Briefly, one million cells were harvested 3 days after transduction and washed twice with cell staining buffer (Biolegend). Cells were resuspended in 100μL of Cell Staining Buffer (Biolegend) and incubated in the dark for 15 minutes at room temperature following the addition of 5μL of Annexin V-APC antibody and 10μL of PI solution, following manufacturer’s instructions. Samples were then analyzed on a BD LSR II (Becton Dickinson, USA) flow cytometer, using BD FACSDiva^TM^ software, and quantified using FlowJo single-cell analysis software (FlowJo, LLC).

### Cell cycle analyses

Cell cycle distribution was analyzed by flow cytometry measurement of cellular DNA content using PI staining. Three million cells were collected, washed, and resuspended in 1 volume of ice-cold 1x Dulbecco’s Phosphate Buffer Saline (PBS, BioConcept) before being fixed by the addition of 2 volumes of ice-cold 100% ethanol during slow vortexing. After 45 min of incubation at 4°C, cells were washed again in ice-cold PBS and resuspended in 1.5 volume of ice-cold PI staining solution (0.1% triton, 200 μg/ml RNAse A, and 50 μg/ml PI). After 30 minutes of incubation at room temperature in the dark, the samples were analyzed on a BD LSR II (Becton Dickinson, USA) flow cytometer, using BD FACSDiva^TM^ software, and quantified using FlowJo single-cell analysis software (FlowJo, LLC).

### CXCL10 measurement in cell culture supernatant

CXCL10 concentration was determined using the Human ProQuantum Immunoassay Kit according to the manufacturer’s instructions. This method relies on antigen recognition by two antibodies, each conjugated to a specific DNA oligo. The presence of the antigen will allow the proximity ligation of the DNA oligo from one antibody to the other, and further quantification of the ligation product by quantitative PCR, as a readout of CXCL10 concentration. Fresh supernatants from U2932 KD and control cells were harvested at day 6 post-transduction. Supernatants were mixed with the two oligonucleotide-conjugated antibodies, diluted 1:10 with assay dilution buffer, and incubated for 1h at room temperature. Oligonucleotide ligation was performed by the addition of Master mix and Ligase and incubation for 1h at room temperature. Quantification of the ligation products was determined by quantitative PCR on an Applied Biosystems 7900HT Fast Real-Time PCT system (ThermoScientific) using the recommended instrument settings. The Ct values were exported to the ProQuantum Cloud application (apps.thermofisher.com/apps/proquantum) to determine CXCL10 concentrations. Finally, supernatant protein concentrations were normalized by cell concentration measured at the beginning of the experiment.

### Analysis of published datasets

The BAM files from 775 exon-enriched DLBCL transcriptomes were downloaded from the European Genome-phenome Archive (EGA) at the European Bioinformatics Institute under the accession number EGAS00001002606^59^. To ensure better analysis quality, we excluded 142 samples with fewer than 1 million counts in total. After filtering lowly expressed genes with an average expression <1 CPM, 23’560 genes were subjected to a univariate Cox regression analysis using the R-package “survival”. For this purpose, each gene expression in normalized counts units was compared to the overall survival (OS) followed by the application of an independent Cox model for each gene. 5’220 genes were found to be significantly associated to survival (Wald test *p-value* <0.05) and 1’530 remained after correction for multiple testing using the Benjamini-Hochberg method (FDR <0.05). KZFP genes were identified using a de novo generated whole genome annotation (Supplementary Table 1) using Hidden Markov Model (HMM) profile generated from seed sequences downloaded from Pfam^96^ and created using hmmbuild from HMMER2 (hmmer.org). KZFP genes were further identified as protein-coding using Ensembl BioMart database (GRCh38.p13, v103)^97^. Other human TF genes were identified using the previously published list from Lambert et al^32^. The categorization of the KZFPs was based on their evolutionary age as previously reported by Imbeault et al^31^.

Clinical data (OS, IPI details, and response to first-line therapy), driver mutation status, and COO classification of DLBCLs were extracted from the metadata provided in the publication of Reddy et al^59^. The relationship between the 41 KZFP genes identified in Cox regression analysis was measured by Spearman rank correlation using the rcorr function of the Hmisc R-package and plotted using pheatmap. *KZFP* genes genomic cluster affiliation and evolutionary ages were extracted from Pontis et al.^43^ and Imbeault et al.^31^, respectively. Hierarchical clustering analysis of lymphoma samples based on the expression of the 18 KZFPs located in the Chr19q co-expression module was computed using default parameters of the heatmap.2 function from gplots using Euclidean distances and Ward’s agglomeration method. The clusterProfiler R-package was used for enrichment analysis of DEGs using Hallmarks gene sets^63^. Copy number alterations were inferred from VCF files provided by Reddy et al. with gains and losses called as described in their publication^59^.

### Data analysis

Unless otherwise specified, graphs were obtained by GraphPad Prism software version 9 or ggplot2 R package.

### RT-qPCR

Total RNA extraction was performed using the NucleoSpin RNA plus kit (Machery-Nagel) according to the manufacturer’s recommendations. cDNA synthesis for qPCR was conducted using the Maxima H minus cDNA synthesis master mix (ThermoScientific). Real-time quantitative PCR was performed using PowerUp SYBR Green Master Mix (ThermoScientific) and run on a QuantStudioTM 6 Flex Real-Time PCR System.

### RNA-seq librairies and downstream analyses

RNA-seq libraries were prepared using the Illumina Truseq Stranded mRNA kit. Libraries were sequenced in 75 or 100 bp paired-end formats on the Illumina Hiseq 4000 and NovaSeq 6000 sequencers, respectively. RNA-seq reads were mapped to the hg19 and T2T-CHM13v2.0 human genome releases using hisat v2.1.0^98^. Only uniquely mapped reads were used for counting over genes and repetitive sequence integrants (MAPQ > 10). Counts for genes and TEs were generated using featureCounts v2 and normalized for sequencing depth using the TMM method implemented in the limma package of Bioconductor^99^. Counts on genes were used as library size to correct both gene and TE expression. For repetitive DNA elements, an in-house curated version of the Repbase database was used. Details about the creation of this list can be found in this reference^71^. Differential gene expression analysis was performed using Voom^100^ as implemented in the Limma package of Bioconductor^101^. P-values were adjusted for multiple testing using the Benjamini-Hochberg method.

### Cleavage Under Targets and Tagmentation (CUT&Tag)

CUT&Tag was performed as described in Kaya-Okur et al^73^ without modifications. For each mark, 150k cells were used per sample using the anti-H3K9me3 primary antibody (Active Motif, AB_2532132), the anti-γH2AX primary antibody (Abcam, ab2893), and anti-rabbit IgG (Abcam, ab46540) secondary antibody. A homemade purified pA-Tn5 protein (3XFlag-pA-Tn5-Fl, addgene #124601) was produced and coupled with MEDS Oligos by the Protein Production and Purification of EPFL, as previously described^102^. Purified recombinant protein was used at a final concentration of 700 ng/uL (1:250 dilution from homemade stock). Libraries were sequenced with 75 bp paired end on the NextSeq 500 (Illumina). Reads were aligned to the T2T-CHM13v2.0 reference genome^74^ using bowtie2^103^. Only proper read pairs with MAPQ>10 were kept. CUT&Tag peaks were called using SEACR v1.3^104^ on “stringent” mode and numeric threshold 0.01 for H3K9me3 and 0.001 for γH2AX, and merged with maximum distance allowed of 2000bp for H3K9me3 and 0bp for γH2AX with bedtools 2.27.1. Encode blacklist for hg19 genome^105,106^ was lifted over to T2T-CHM13v2.0 genome and filtered out from the annotated peaks. Bedtools multicov was used to count mapped reads on the annotated and filtered peaks, and differential peak analysis was performed using Voom after library size correction (using the total number of aligned reads as size factor) performed using the TMM method.

Bigwig coverage tracks with the mean of replicate samples were generated using bedtools 2.27.1^107^ and deeptools 3.3.1^108^, and heatmap representations of the coverage signal were performed using computeMatrix function and plotHeatmap from deeptools 3.3.1.

### EdU DNA synthesis monitoring flow cytometry

Lymphoma cells were pulse-labeled with 10 uM 5-ethynyl-2′-deoxyuridine (EdU) for 20 min and subsequently fixed with 2% formaldehyde for 30 min at room temperature (RT). EdU incorporation was detected using Click chemistry according to manufacturer’s instructions (Click-iT EdU Flow Cytometry Cell Proliferation Assay, Invitrogen). Cells were resuspended in 1x Phosphate Buffer Solution (PBS, BioConcept) with 1% bovine serum albumin (BSA), 2 ug/ml DAPI and 0.5 mg/ml RNase A for 30 min at RT and subsequently analyzed on a BD LSR II (Becton Dickinson, USA) flow cytometer, using BD FACSDivaTM software, and quantified using FlowJo single-cell analysis software (FlowJo, LLC).

### TrAEL-Seq

TrAEL-Seq was performed as described in the publication of Kara et al^109^. In Brief, 1 million of cells transduced with ShScr and ShRNA.1 were collected 3 days after LV transduction. Cells were washed once in 10 ml L buffer (10 mM Tris HCl (pH 7.5), 100 mM EDTA, 20 mM NaCl) and resuspended in 60 μl L buffer in a 2-ml tube. Samples were heated to 50°C for 2 to 3 min before the addition of 40 μl molten CleanCut agarose (Bio-Rad 1703594), vortexed vigorously for 5 sec before pipetting in plug mould (Bio-Rad 1703713), and solidified at 4°C for 30 min. Each plug was transferred to a 2-ml tube containing 500 μl digestion buffer (10 mM Tris HCl (pH 7.5), 100 mM EDTA, 20 mM NaCl, 1% sodium N-lauroyl sarcosine, 0.1 mg/ml Proteinase K) and incubated overnight at 50°C. Plugs were washed and treated with RNase T1. Half of an agarose plug was used for each library. For restriction enzyme digestion, a plug was equilibrated 30 min in 200 μl 1x CutSmart buffer (NEB), digested overnight at 37°C with 1 μl 20 U/μl NotI-HF (NEB R3189S) and 1 μl 10 U/μl PmeI (NEB R0560S) in 400 μl 1x CutSmart buffer, then 1 μl 20 U/μl SfiI (NEB R0123S) was added and incubation continued overnight at 50°C. The plug was rinsed with 1x TE before further processing. Plugs were equilibrated once in 100 μl 1x TdT buffer (NEB) for 30 min at room temperature, then incubated for 2 h at 37°C in 100 μl 1x TdT buffer containing 4 μl 10 mM ATP and 1 μl Terminal Transferase (NEB M0315L). Plugs were rinsed with 1 ml Tris buffer (10 mM Tris HCl (pH 8.0)), equilibrated in 100 μl 1x T4 RNA ligase buffer (NEB) containing 40 μl 50% PEG 8000 for 1 h at room temperature, then incubated overnight at 25°C in 100 μl 1x T4 RNA ligase buffer (NEB) containing 40 μl 50% PEG 8000, 1 μl 10 pM/μl and previously adenylated using the 5′ DNA adenylation kit (NEB, E2610S), and 1 μl T4 RNA ligase 2 truncated KQ (NEB M0373L). Plugs were then rinsed with 1 ml Tris buffer, transferred to 15 ml tubes, and washed 3 times in 10 ml Tris buffer with rocking at room temperature for 1 to 2 h each, then washed again overnight under the same conditions. Plugs were equilibrated for 15 min with 1 ml agarase buffer (10 mM Bis-Tris-HCl, 1 mM EDTA (pH 6.5)), then the supernatant removed and 50 μl agarase buffer added. Plugs were melted for 20 min at 65°C, transferred for 5 min to a heating block preheated to 42°C, 1 μl β-agarase (NEB M0392S) was added and mixed by flicking without allowing sample to cool, and incubation continued at 42°C for 1 h. DNA was ethanol precipitated with 25 μl 10 M NH4OAc, 1 μl GlycoBlue, 330 μl of ethanol and resuspended in 10 μl 0.1x TE. A volume of 40 μl reaction mix containing 5 μl isothermal amplification buffer (NEB), 3 μl 100 mM MgSO4, 2 μl 10 mM dNTPs, and 1 μl Bst 2 WarmStart DNA polymerase (NEB M0538S) was added and sample incubated 30 min at 65°C before precipitation with 12.5 μl 10 M NH4OAc, 1 μl GlycoBlue, 160 μl ethanol and redissolving pellet in 130 μl 1x TE. The DNA was transferred to an AFA microTUBE (Covaris 520045) and fragmented in a Covaris E220 using duty factor 10, PIP 175, Cycles 200, Temp 11°C, then transferred to a 1.5-ml tube containing 8 μl prewashed Dynabeads MyOne streptavidin C1 beads (Thermo, 65001) resuspended in 300 μl 2x TN (10 mM Tris (pH 8), 2 M NaCl) along with 170 μl water (total volume 600 μl) and incubated 30 min at room temperature on a rotating wheel. Beads were washed once with 500 μl 5 mM Tris (pH 8), 0.5 mM EDTA, 1 M NaCl, 5 min on wheel and once with 500 μl 0.1x TE, 5 min on wheel before resuspension in 25 μl 0.1x TE. TrAEL-seq adaptor 2 was added using a modified NEBNext Ultra II DNA kit (NEB E7645S): 3.5 μl NEBNext Ultra II End Prep buffer, 1 μl 1 ng/μl sonicated salmon sperm DNA (this is used as a carrier), and 1.5 μl NEBNext Ultra II End Prep enzyme were added and reaction incubated 30 min at room temperature and 30 min at 65°C. After cooling, 1.25 μl 10 pM/μl TrAEL-seq adaptor 2, 0.5 μl NEBNext ligation enhancer, and 15 μl NEBNext Ultra II ligation mix were added and incubated 30 min at room temperature. The reaction mix was removed and discarded and beads were rinsed with 500 μl wash buffer (5 mM Tris (pH 8), 0.5 mM EDTA, 1 M NaCl), then washed twice with 1 ml wash buffer for 10 min on wheel at room temperature and once for 10 min with 1 ml 0.1x TE. Libraries were eluted from beads with 11 μl 1x TE and 1.5 μl USER enzyme (NEB) for 15 min at 37°C, then again with 10.5 μl 1x TE and 1.5 μl USER enzyme (NEB) for 15 min at 37°C, and the 2 eluates combined. Amplification of the library was performed with components of the NEBNext Ultra II DNA kit (NEB E7645S) and a NEBNext Multiplex Oligos set (e.g., NEB E7335S). An initial test amplification was used to determine the optimal cycle number for each library. For this, 1.25 μl library was amplified in 10 μl total volume with 0.4 μl each of the NEBNext Universal and any NEBNext Index primers with 5 μl NEBNext Ultra II Q5 PCR master mix. Cycling program: 98°C 30 s, then 18 cycles of (98°C 10 s, 65°C 75 s), 65°C 5 min. Test PCR was cleaned with 8 μl AMPure XP beads (Beckman A63881) and eluted with 2.5 μl 0.1x TE, of which 1 μl was examined on a Bioanalyser high sensitivity DNA chip (Agilent 5067–4626). Ideal cycle number should bring final library to final concentration of 1 to 3 nM, noting that the final library will be 2 to 3 cycles more concentrated than the test anyway. A volume of 21 μl of library was then amplified with 2 μl each of NEBNext Universal and chosen Index primer and 25 μl NEBNext Ultra II Q5 PCR master mix using same conditions as above for calculated cycle number. Amplified library was cleaned with 40 μl AMPure XP beads (Beckman A63881) and eluted with 26 μl 0.1x TE, then 25 μl of this was again purified with 20 μl AMPure XP beads and eluted with 11 μl 0.1x TE. Final libraries were quality controlled and quantified using a Bioanalyzer system (2100 Agilent 5067–4626) and KAPA qPCR (Roche KK4835).

Preparation of TrAEL-seq adaptors: DNA oligonucleotides were synthesized and PAGE purified by Sigma-Genosys (Merck, United Kingdom). Sequences were as follows:

1. [Phos]NNNNNNNNAGATCGGAAGAGCGTCGTGTAGGGAAAGAGTGTUGCGCAGGCCATTGGCC[BtndT] GCGCUACACTCTTTCCCTACACGACGCT.
2. [Phos]GATCGGAAGAGCACACGTCTGAACTCCAGTCUUUUGACTGGAGTTCAGACGTGTGCTCTTCCGA TC*T.

Libraries were sequenced on an Illumina NextSeq 500 sequencer as 75 bp paired-end reads. All scripts used for the analyses of TrAEL-seq data can be found in the following git repository: https://github.com/fmartins/traelseq). The paired-end TrAEL-seq reads carry an 8-bp in-line barcode (UMI) at the 5′-end of the first read in the pair, followed by a variable number of 1 to 3 thymines (T). Read structure of the first read in the pair is, therefore, NNNNNNNN(T)nSEQUENCESPECIFIC, where NNNNNNNN is the UMI, and(T)n is the poly(T). We used flexbar^110^ to 1) remove the first 8 bp (UMI) of each read, 2) for each read in the pair, add the UMI sequence to the end of the readID, and 3) remove up to 3 T (inclusive) at the start of the sequence. UMI processed files were then aligned to the T2T-CHM13v2.0 genome using Bowtie2^103^ (parameters: --local -D 15 -R 2 -N 1 -L 25 -i S,1,0.75). Alignment files (bams) were then deduplicated using umit_tools^111^ (parameters: --paired --extract-umi-method read_id --method unique). Reads were then summed in running windows using the bamCoverage function from deepTools^108^. Count tables per region in the format of bedgraph files were then imported into R^112^ for plotting and statistics. Bigwig files were computed using the bedGraphToBigWig software^113^. For read polarity plots, the following formula was used: read polarity = (R − F)/(R + F), where F and R relate to the total forward and reverse read counts respectively. To compute R and F, the following parameters were used in bamCoverage: for forward (--samFlagInclude 64 --samFlagExclude 16) and for reverse (--samFlagInclude 80).

### Immunofluorescence

U2932 cells were fixed and permeabilized using True-Nuclear^TM^ Transcription Factor Buffer Set (Biolegend) following manufacturer’s instructions, incubated with primary anti-dsDNA antibody (1:100, Mybiosource), anti-phospho-STING Ser365 (1:400, CST), or anti-phospho-IRF3 Ser396 (1:100, CST) for 30 minutes, washed, and incubated with secondary anti-Mouse IgG Alexa Fluor 488 (1:100, Invitrogen) or anti-Rabbit IgG Alexa Fluor 568 (1:100, Invitrogen) for another 30 minutes. Nuclei were stained with <u>DAPI</u> (4′, 6-diamidino-2-phenylindole, 1 μg/mL, Sigma-Aldrich). Images were acquired on a point-scanning confocal microscope SP8 Leica with an oil 40x objective 1.25 N.A. Images are Z-stacks of dimensions 256×256 in XY and spanning the entire cell in Z (voxel size 120×120×500 nm). Pinhole was set at 1 Airy Unit for an emission wavelength of 520 nm for all channels, acquired in sequential scan mode. DAPI was excited with a 405 nm laser (emission 420-470nm), its emission collected between 420 and 470 nm. The excitation of the DAPI channel is used to make a transmission image. DsDNA (Alexa 488) was excited with a 488 nm laser, its emission collected between 500 and 530nm. Phospho-STING and phospho-IRF3 (Alexa 568) were excited with a 552 nm laser, their emission was collected between 560 and 610 nm. DsDNA quantification estimation was performed in ImageJ/Fiji^114^. Each image is projected in Z using maximum intensity projection. The DAPI channel was subsequently median filtered (radius = 2 pixels), and the nuclear region was delineated by using a fixed intensity threshold, identical for all images. A band of 0.8 micrometer surrounding the nucleus is then defined by shrinking the nucleus region by 5 pixels, then expanding it by 0.8 micrometer, thus defining a cytosolic band region. Average intensities of DAPI and dsDNA channels were then measured within the shrank nuclear region and the cytosolic band. All experiments were reproduced at least once.

### Phagocytosis assay

Human M1 macrophages (effector cells) were plated into 96-well flat clear bottom black walled culture plates (ThermoScientific 165305) at a density of 1×10^4^- 2.5×10^4^ cells/well in 50 uL complete RPMI medium supplemented with M-CSF (50 ng/ml) overnight. U2932 cells (target cells) were transduced as described above, harvested at day 4, washed with pHrodo Wash bBffer (Sartorius), and labeled using pHrodo red (Sartorius) at a final concentration of 31.3 ng/mL for 45 minutes at 37°C in pHrodo Labeling Buffer (Sartorius), before being washed again and resuspended in complete RPMI medium. 3×10^4^-7.5×10^4^ target cells/well were then seeded over effector cells at target:effector ratio of 3:1. 96-well plates were then placed into the InCucyte S3 System (Sartorius) for time-lapse imaging. The pHrodo signal was measured by the acquisition of images from 3-5 fields/well every 1 to 1:30 hours for 24 hours.

### HLA typing and prediction of HLA binding affinity

HLA typing using HLA-HD v1.4.0^115^ on RNA sequencing data from U2932. For each sample, the consensus HLA haplotypes were determined as the consensus typing derived from all RNA sequencing replicates. NetMHCpan-4.1 tool was used for predicting the binding affinity of the eluted peptides to the respective HLA alleles^116^.

### Immunopeptidomics analysis

HLA-I peptides were immunopurified from U2932 cells. A first set of control and *ZNF587/417* KD cell pellets (30 million cells each) was used to generate a spectral library from data-dependent and data-independent acquisition methods (DDA and DIA, respectively). A second set of pellets (with at least 20 millions of cells) were acquired using DIA method and analyzed by Spectronaut. The two sets of samples were processed as previously described to purify the HLA-I peptides^117^. Briefly, anti-pan HLA-I monoclonal antibodies (W6/32) were purified from supematant of HB95 hybridoma cells (ATCC® HB-95) using protein-A sepharose 4B (Pro-A) beads (Invitrogen) and cross-linked to pro-A beads. Each sample was lysed in 1.5mL lysis buffer comprising of PBS containing 0.25% sodium deoxycholate (Sigma-Aldrich), 0.2 mM iodoacetamide (Sigma-Aldrich), 1 mM EDTA, 1:200 Protease Inhibitors Cocktail (Sigma-Aldrich), 1 mM Phenylmethylsulfonylfluoride (Roche, Basel, Switzerland), 1% octyl-beta-D glucopyranoside (Sigma-Alrich), at 4°C for 1 hour and centrifuged at 4°C at 14200 rpm for 50 min using a refrigerated table-top centrifuge (Eppendorf Centrifuge, Hamburg, Germany). The cleared lysates were loaded on wells of 96-well single-use filter micro-plates with 3 µm glass fibers and 25 µm polyethylene membranes (Agilent, 204495-100) containing the HB95 cross-linked beads. Once the lysates flew through, the plates were washed several times with solutions containing various concentrations of salts and two final washes of 2mL of 20mM Tris-HCl pH8 using the Waters Positive Pressure-96 Processor (Waters). The HLA complexes bound to the beads were then directly eluted into preconditioned Sep-Pak tC18 100 mg Sorbent 96-well plates (Waters, 186002321) using 1% trifluoroacetic acid (TFA; Sigma Aldrich). After two washes of 1mL 0.1% TFA, the HLA-I peptides were eluted using 25% acetonitrile (ACN; Sigma Aldrich) in 0.1% TFA. Peptides were dried using vacuum centrifugation (Concentrator plus, Eppendorf) and stored at −20 °C. Priorr to LC-MS/MS analysis, iRT peptides (Biognosys) were spiked in each sample for retention time calibration through Spectronaut analysis. For each sample, dried peptides were re-suspend in 12 ul of which 3ul samples were injected for DDA and 2 ul for DIA.

The Easy-nLC 1200 liquid chromatograph (Thermo Fisher Scientific) was coupled to a Q Exactive HF-X mass spectrometer (Thermo Fisher Scientific) for peptide tandem mass spectrometry data analysis. Peptides were separated on a home-made 50cm analytical column with a tip size of ∼10um (OD = 365um, ID = 75um) and packed with ReproSil-Pur C18 (1.9-μm particles, 120 Å pore size, Dr Maisch GmbH). The peptide separation by chromatography was performed as previously described^118^. For MS/MS data acquisition in DDA and DIA, the same ion sampling methods described by Pak et al.^119^ were used.

### DDA and DIA data analysis

We used Spectronaut 15.7.220308.50606^120^ to build a spectral library from DDA and DIA MS/M data. The peptide identification search was done by Pulsar^TM^ using an unspecific digestion type and a peptide length from 8 -15 a.a. with a maximum variable modification of 5 per peptide. Acetylation of n-term and oxidation of methionine were used as variable modifications. The library was then generated using a peptide and PSM FDR ≤ 0.01 and protein FDR = 1 from Pulsar search. The DIA MS/MS data was matched against the library as previously described by Pak et al.^119^, with the exception that this time iRT peptide calibration was used.

### Data Availability

The accession number for the RNA-seq and CUT&Tag data generated in this article is GEO:

## ACKNOWLEDGMENTS

We thank C. Raclot, P.-Y. Helleboid, J. Duc, V. Glutz, R. Guiet, N. Chiaruttini and J. Pontis for their technical support and scientific advice; R. Guindon for help in graphical output of the figures, the EPFL Flow Cytometry (FCCF), Protein Production and Purification (PTPSP) and Gene Expression (GECF) core facilities for the use of their systems and services. We thank A. Mayran and the Laboratory of Developmental Genetics of Prof. Denis Duboule for help with the use of their InCucyte S3 Sartorius imaging system. We also acknowledge all the members of the Trono laboratory for fruitful discussions and constant support. This study was supported by grants from the Swiss National Science Foundation and the European Research Council (KRABnKAP, no. 268721; Transpos-X, no. 694658), the Personalized Health and Related Technologies (PHRT) strategic focus area of the Swiss Federal Institutes of Technology (ETH) Domain and the Swiss Personalized Health Network (SPHN) initiative of the Swiss Academy of Medical Sciences (project no. 2017-407) and by the generous longstanding support of the Aclon Foundation to D.T.

## AUTHOR CONTRIBUTIONS

F.M. and D.T. conceived the study and designed experiments; F.M., O.R., and R.F. performed the wet experiments with the technical support of S.O.; J.C., E.P., C.P., and F.M. conducted the bioinformatics analyses, and F.M., O.R., R.F., J.CF., and D.T. wrote the manuscript, with review and corrections by all authors.

## CONTACT FOR REAGENT AND RESOURCE SHARING

Further information and requests for resources and reagents should be directed to the lead contact, Didier Trono.

---

**Figure S1 related to Figure 2:**
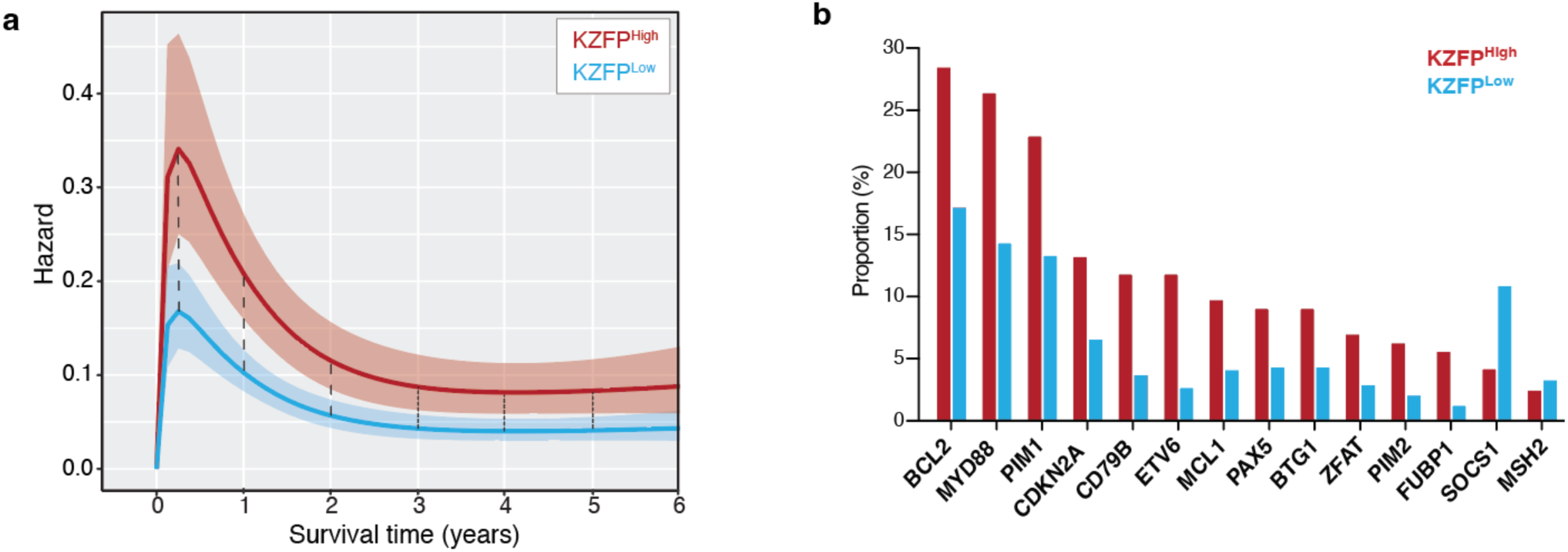
A module of KZFP genes in DLBCLs with cell-autonomous growth features. (a) Smoothed hazard functions by KZFP^High/Low^ grouping for death among patients treated with a standard rituximab-based regimen, with, in dark red the hazards of patients diagnosed with a KZFP^High^ and in blue with a KZFP^Low^ DLBCL. Vertical dashed lines indicate the difference in hazard between the two groups at given time points after diagnosis. (b) Bar plots depicting the proportion of samples with given driver mutations in KZFP^High^ (dark red) and KZFP^Low^ (blue) groups. Only significantly enriched or depleted (FDR <0.05) driver mutations are depicted. Statistics: Two-tailed Fisher’s Exact Test.

**Figure S2 related to Figure 3:**
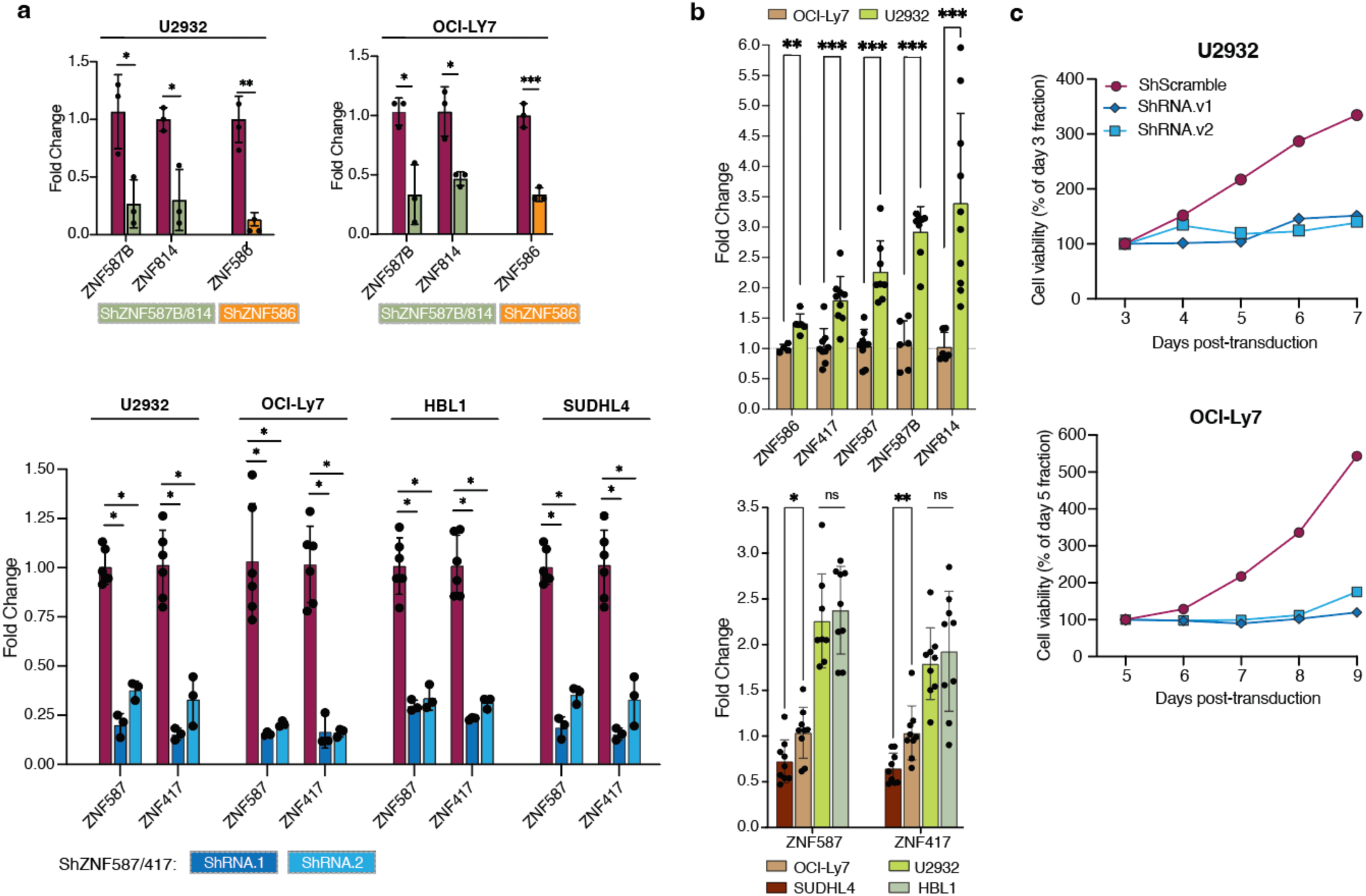
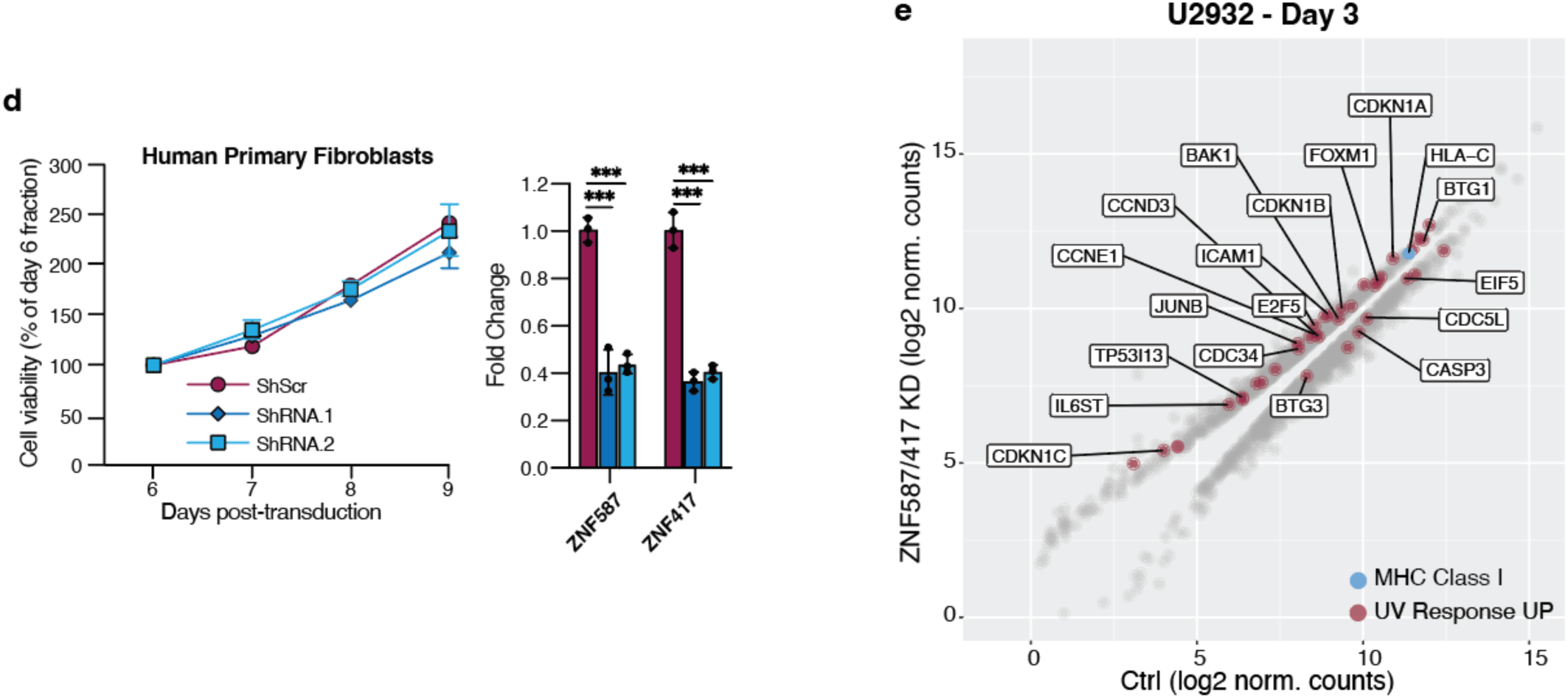
ZNF587/417 depletion impairs lymphoma cell growth and viability. (a) Quantitative RT-PCR analysis of ZNF587B, ZNF814, ZNF587, ZNF417, and ZNF586 mRNAs 5 days after LV transduction with shRNAs targeting ZNF586, ZNF587B/ZNF814, and ZNF587/ZNF417 paralog pairs. Statistics: Student’s t-test. (b) Top: Bar plots showing the relative expression of ZNF586, ZNF417, ZNF587, ZNF587B, and ZNF814 in U2932 compared to OCI-Ly7 using GUSB, TBP, and/or ALAS2 housekeeping genes for normalization. Bottom: Bar plots showing the relative expression of ZNF587 and ZNF417 in SUDHL4, U2932, HBL1 cells compared to OCI-Ly7 using GUSB, TBP, and/or ALAS2 housekeeping genes for normalization. (c) MTT proliferation assays of U2932 and OCI-Ly7 upon LV transduction with two different anti-ZNF587/417 shRNAs, or control shRNA (shScr). Cells were plated after 5 days of LV transduction and 3 days of puromycin selection. (d) MTT proliferation assay of human primary dermal fibroblasts upon LV transduction with two anti-ZNF587/417 shRNAs or control shRNA. Cells were plated after 5 days of LV transduction and 3 days of puromycin selection. Quantitative RT-PCR analysis of ZNF587 and ZNF417 mRNAs 5 days after LV transduction. Statistics: Student’s t-test. (e) Scatter plot of RNA-seq from ZNF417/587 U2932 KD versus shScr control cells at day 3 of KD, outlining DEGs (grey dots, FDR <0.05) and among them genes belonging to the UV response UP Hallmark gene set (dark red dots) and MHC Class I genes (blue dots).

**Figure S3 related to Figure 4:**
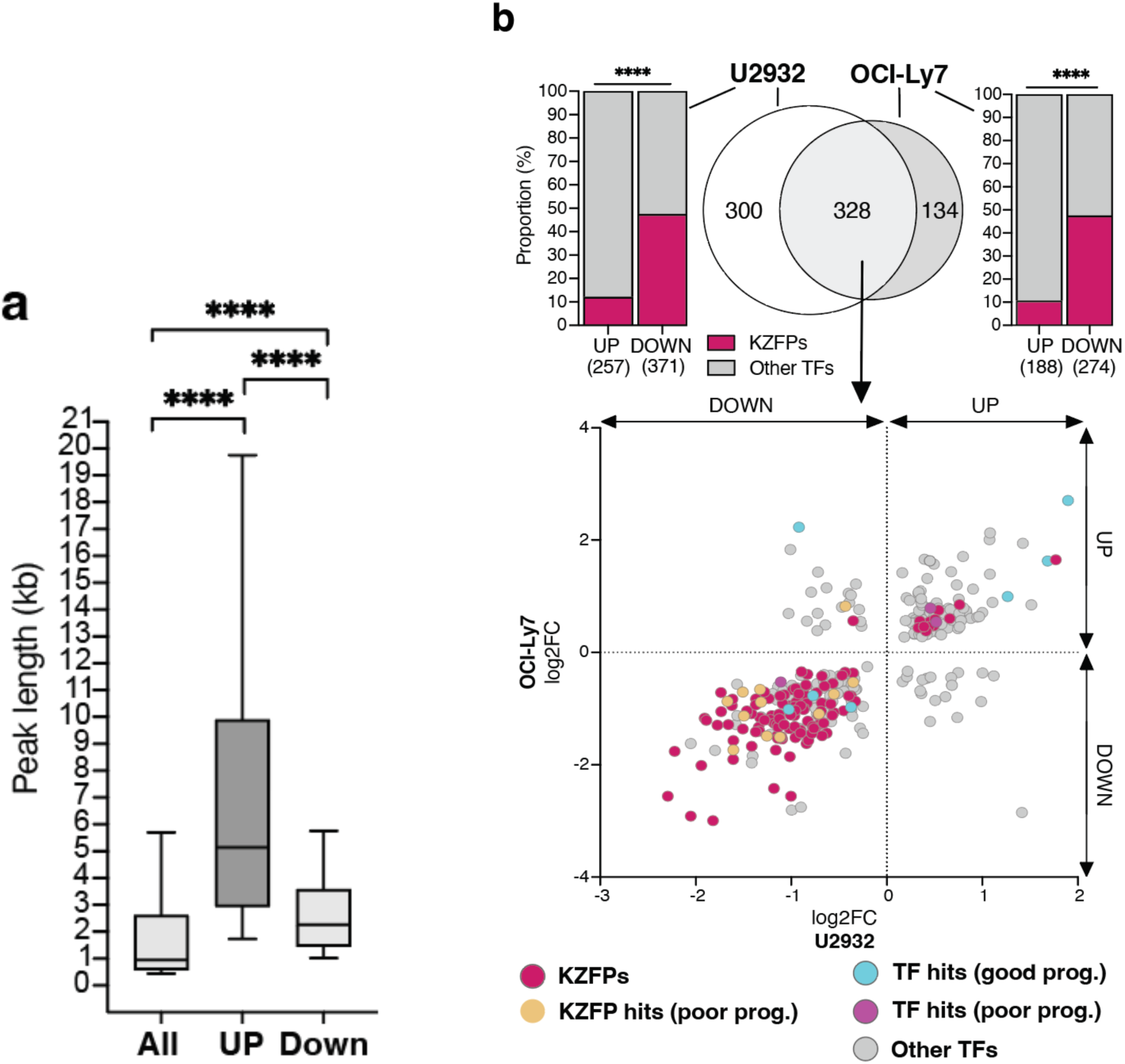
ZNF587/417 depletion alters the heterochromatin landscape of lymphoma cells. (a) Box plots showing the quartiles, the 10^th^ and 90^th^ percentiles (whiskers) of all, gaining, and losing H3k9me3 peaks lengths (in kilo-bases). Statistics: Two-sided Mann–Whitney U-test. (b) Top: Venn diagrams of differentially expressed TFs (FDR <0.05) upon ZNF587/417 KD in U2932 and OCI-Ly7 cells 6 days after LV transduction. The number of differentially expressed TFs shared between the two cell lines is shown in the overlapping area of the two diagrams. The number of TFs differentially expressed in only one of the two cell lines is shown in the non-overlapping area of the diagrams. Stacked bar charts depicting the proportion of up- and downregulated KZFPs amongst up- and downregulated TFs in U2932 and OCI-Ly7 are shown on each side of the Venn diagrams Bottom: Scatterplot of log2 fold changes of DE TFs shared between U2932 (x-axis) and OCI-Ly7 (y-axis) KD cells. Different colors were used to highlight TFs and KZFPs that were identified as positive hits from the Cox regression analysis screen reported in Figure 1.

**Fig. S4 related to Figure 5:**
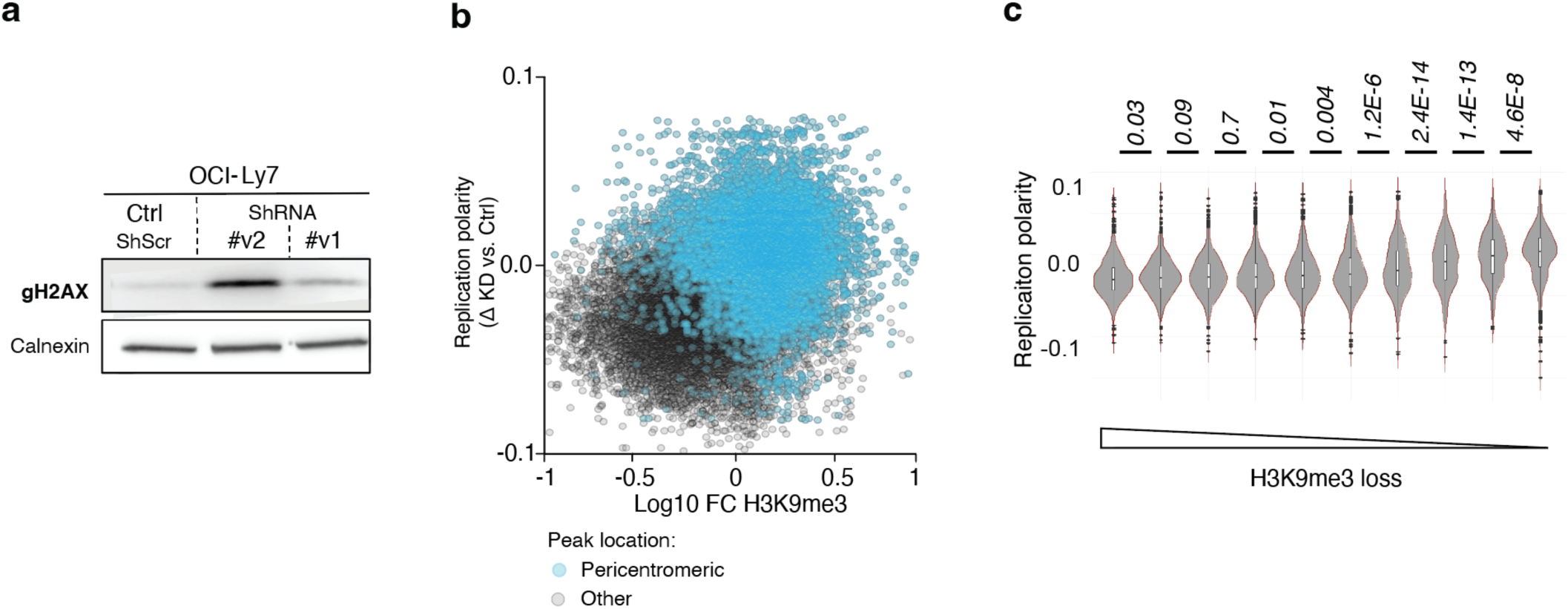
ZNF587/417 depletion triggers replicative stress in lymphoma cells. (a) Western blot analysis of *γ*H2AX in OCI-Ly7 cells 3 days after LV transduction with 2 different anti-ZNF587/417 shRNAs and control shRNA (shScr). Calnexin was used as a loading control. (b) Scatter plot of replication polarity in function of H3K9me3 changes idem as Figure 5h. H3K9me3 peaks in pericentromeric regions are highlighted in blue and the ones in other regions in grey. (c) Violin plots of replication polarity deltas between KD and control conditions calculated as in Figure 5h grouped in 10 bins of the same numbers of H3K9me3 peaks and ranked by decreasing H3K9me3 loss. Statistics: Two-sided Mann–Whitney U-test.

**Figure S5 related to Figure 6:**
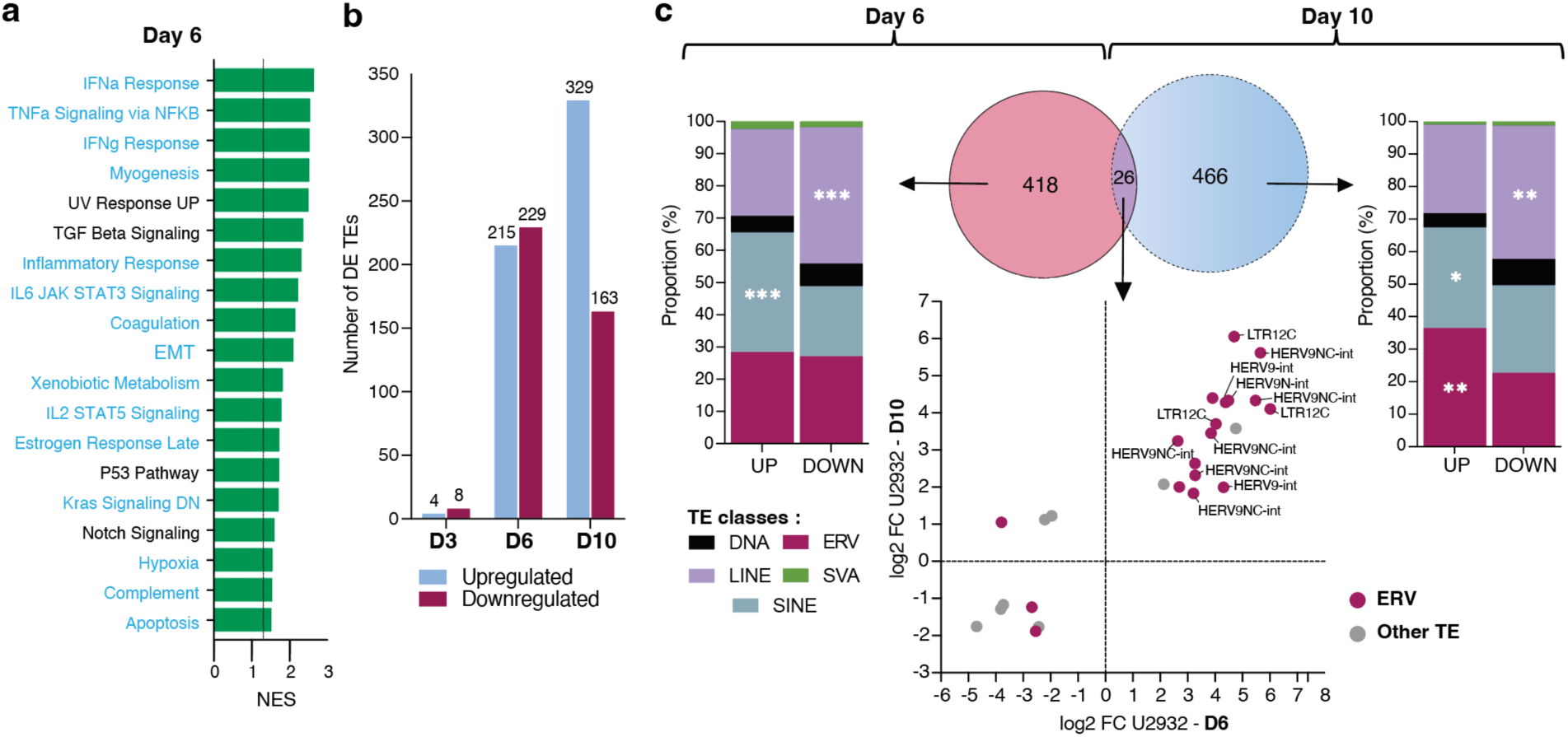
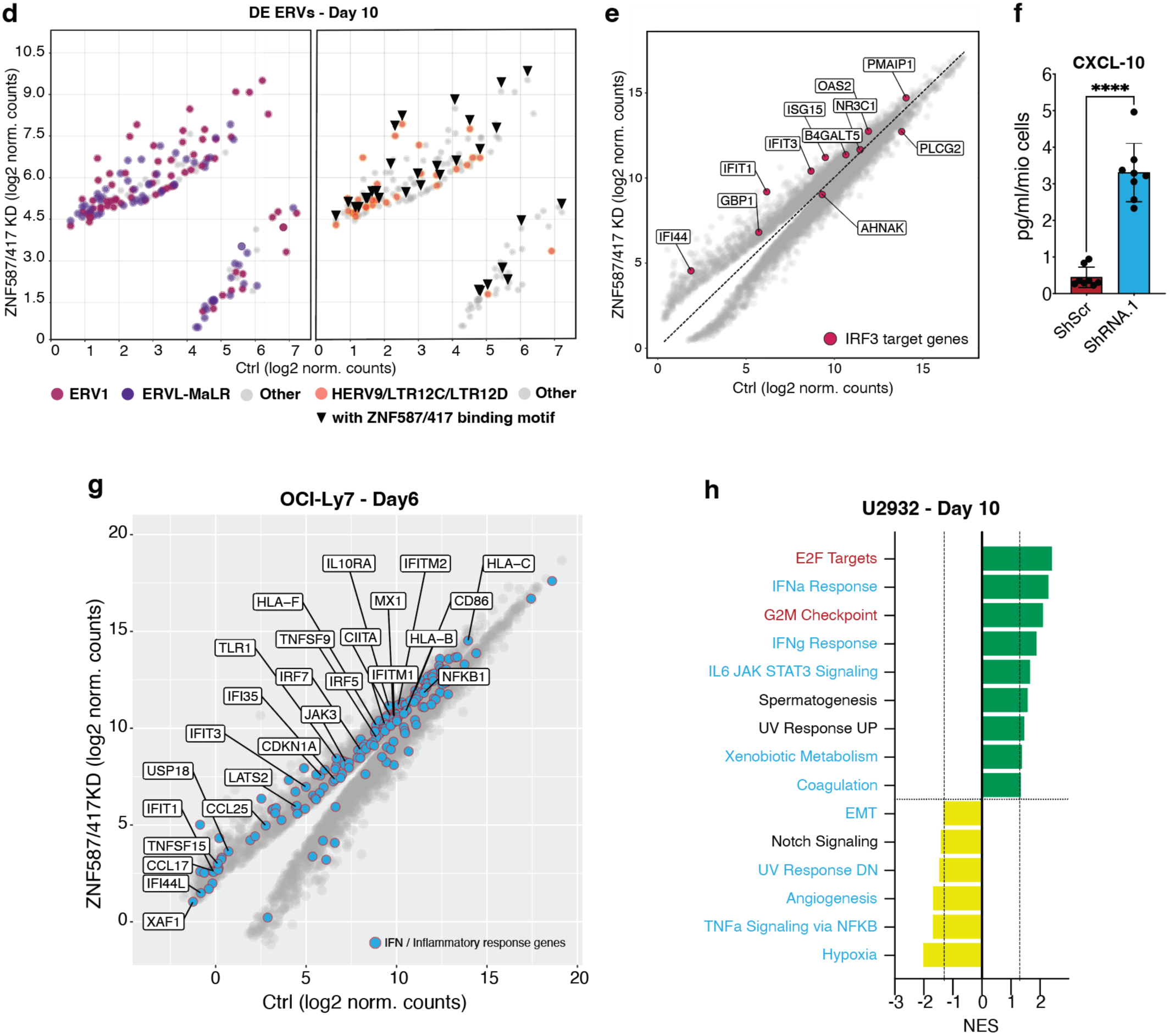
ZNF587/417 depletion leads to cell-intrinsic inflammation. (a) Waterfall plot of Hallmark GSEA signatures ranked by NES issued from RNA-seq data of day 6 ZNF587/417 KD in U2932 cells. Enriched signatures (NES>0) are highlighted in green. No signature was found significantly depleted. The dotted line represents p-value cutoff <0.05. Hallmark labels found enriched in KZFP^Low^ from Fig. 2 are highlighted in blue. (b) Bar plot depicting the number of DE TE integrants (FDR <0.05, FC >2) at day 3, 6, and 10 of ZNF587/417 KD in U2932 cells. Control shRNA-transduced cells collected in parallel to shRNA.1 KD cells were used as controls for every time point. (c) Center: Venn diagrams of DE TEs (FDR <0.05) upon ZNF587/417 KD in U2932 cells at day 6 (left) and 10 (right) after LV transduction. The number of DE TEs shared between the two time-points is shown at the intersection of the two disks. The number of TEs differentially expressed in only one of the two cell lines is shown in the non-overlapping area of the disks. Stacked bar charts depicting the proportion of up- and downregulated TEs at day 6 (left) and 10 (right) in U2932 KD cells are shown on each side of the Venn diagrams Bottom: Scatterplot of log2 fold changes of DE TEs shared between day 6 (x-axis) and day 10 (y-axis) in U2932 KD cells (ERVs were highlighted in dark red and other TEs in gray dots). (d) Scatterplot of log2 fold changes of DE ERVs at day 10 in U2932 KD cells. ERV1 elements are highlighted with dark red dots, ERVL-MaLR with purple dots, and other ERVs with gray dots. ERVs with a ZNF587 and/or ZNF417 binding motif are pointed with a triangle. The top 3 binding motifs determined in H1 embryonic stem cells were used^71^ for both KZFPs. (e) Scatter plot of RNA-seq from ZNF417/587 U2932 KD versus (shScr) control cells at day 6 of KD, outlining DEGs (grey dots, FDR <0.05) and among them the genes reported as IRF3 targets by Grandvaux et al.^121^ (dark red dots). (f) Bar plot measuring the concentration of CXCL10 cytokine by immunoassay in cell culture supernatant of ZNF587/417 shRNA.1 KD (blue) and control cells (dark red) 6 days after LV transduction normalized by the number of cells in each condition (pg/ml/millions of cells). (g) Scatter plot of RNA-seq from ZNF417/587 OCI-Ly7 KD versus control cells at day 6 of KD, outlining DEGs (grey dots, FDR <0.05) and among them genes belonging to type I/II IFN and Inflammatory response Hallmark gene sets, Interferon Signaling Reactome, and cellular response to type I, II and III IFN gene ontology terms (blue dots). (h) Waterfall plot of Hallmark GSEA signatures ranked by NES issued from RNA-seq data of day 10 ZNF587/417 KD in U2932 cells. Enriched signatures (NES>0) are highlighted in green and depleted signatures (NES<0) in yellow. The dotted line represents p-value cutoff <0.05. Hallmark labels found enriched in KZFP-Low/-High groups from Fig. 2 are highlighted in blue and dark red, respectively.

